# Behavioral adaptation to changing energy constraints via altered frequency of movement selection

**DOI:** 10.1101/2023.11.08.566262

**Authors:** Thomas Darveniza, Shuyu I. Zhu, Zac Pujic, Biao Sun, Matthew Levendosky, Robert Wong, Ramesh Agarwal, Michael H. McCullough, Geoffrey J. Goodhill

**Affiliations:** Departments of Developmental Biology and Neuroscience, Washington University in St. Louis; Department of Electrical and Systems Engineering, Washington University in St. Louis; Department of Mechanical Engineering and Materials Science, Washington University in St Louis; Queensland Brain Institute, The University of Queensland, Brisbane, Queensland 4072, Australia; School of Mathematics and Physics, The University of Queensland, Brisbane, Queensland 4072, Australia; College of Engineering, Computing and Cybernetics, Australian National University

## Abstract

Animal behavior is strongly constrained by energy consumption. A natural manipulation which provides insight into this constraint is development, where an animal must adapt its movement to a changing energy landscape as its body grows. Unlike many other animals, for fish it is relatively easy to estimate the energy consumed by their movements via fluid mechanics. Here we simulated the fluid mechanics of *>*100,000 experimentally-recorded movement bouts from larval zebrafish across different ages and fluid conditions as they hunted *Paramecia*. We find that these fish adapt to their changing relationship with the fluid environment as they grow by adjusting the frequency with which they select different types of movements, so that more expensive movements are chosen less often. This strategy was preserved when fish were raised in an unnaturally viscous environment. This work suggests a general principle by which animals could minimize energy consumption in the face of changing energy costs over development.

## Introduction

Our understanding of unconstrained animal behavior has increased dramatically in recent years. The introduction of powerful new techniques from computer vision for automated tracking of animal pose has allowed much larger datasets to be generated than was possible using manual tracking [1, 2]. With these new datasets, a variety of unsupervised machine-learning methods can now automatically segment these behaviors into behavioral syllables and allow underlying patterns in behavioral sequences to be revealed [3–5]. This in turn is enabling an understanding of how neural circuits drive these behaviors [5–8].

However although these animals are freely moving, their behavior is still subject to numerous constraints. The most fundamental of these is energy consumption: the energetic costs of behaviors must be less than the rate of energy acquisition that is achieved by these behaviors [9]. There is thus strong pressure for behaviors to be energetically efficient, and continually adapt to changes in efficiency caused by ongoing bodily or environmental changes. How neural circuits monitor energy efficiency and adjust behaviors in light of changing energetic constraints is unknown.

Unique opportunities for analysis of energetics are offered by fish [10]. Fish movement patterns are simpler to describe than multi-limbed organisms such as primates, rodents and flies, and their interactions with the fluid environment can be well understood via the Navier-Stokes equations. A particularly interesting case is that of larval fish, where the interactions experienced between body morphology, movement and the environment can change rapidly over development [11]. Such developmental changes act as a natural system perturbation, providing an attractive model for testing the influence of energetic constraints on behaviour.

Here we focus on larval zebrafish, since it is a well-understood model for studying both development [12–17] and neural circuit control of behavior [18–43]. Between 3 and 5 days post-fertilization (dpf) zebrafish transition from cyclical to burst-and-coast locomotion [44, 45]. This is likely because cyclical swimming is more efficient in viscous regimes where drag is high (low Reynolds number; this is a dimensionless quantity which measures the ratio of inertial to viscous forces on a body of given size and speed in a fluid), while burst-and-coast swimming becomes more efficient as the Reynolds number increases [46]. However by 5 dpf zebrafish have depleted the energy reserves contained in their yolk sac and begin to actively acquire energy by hunting moving prey such as *Paramecia* [47]. How changes in energetics caused by their subsequent growth constrain behavioral algorithms for hunting is unknown.

Previous work has shown that escape bouts in response to predatory cues match predictions based on optimising distance subject to energy constraints [48–50]. Similarly rheotactic behaviors, the automatic response of fish to maintain position in flowing water, use a duty cycle that minimises the Cost of Transport (COT) [51]. However such analyses have often used idealized forms of movement, and it is not clear how closely these replicate natural movements. Furthermore hunting behaviors, one of the most biologically important behaviors of predatory larval fish, have not previously been analyzed in terms of energetics. While hunting sequences have been hypothesised to minimise energy cost [52], how this might be implemented algorithmically is unclear. In addition, previously-proposed optimisation principles have yet to be tested across the differing fluid regimes that occur during development [53], and thus it is unknown whether these principles form part of a generalised computational strategy.

Here, we first construct the behavioral space of larval zebrafish over multiple developmental stages while they hunt prey. We find that the basic structure of movement bouts is preserved over development, but their selection probabilities change. We then introduce a method for computational fluid dynamics (CFD) simulation of experimentally-recorded larval zebrafish movement bouts using the Immersed Boundary Adaptive Mesh Refinement (IBAMR) infrastructure [54], allowing for energy consumption estimations for swimming motions. This reveals that the relative ordering of energy consumption across different bout types changes over development. Remarkably however the probability of bout selection for hunting behaviors adapts over development to maintain a close-to-monotonic relationship with the energy required for each bout. We then confirm the robustness of this finding by performing a similar analysis of larval zebrafish reared in a high-viscosity environment. Together this work suggests how changing energy landscapes could influence and constrain emergent patterns of complex behaviors over development.

## Results

### Bout types are stable over development but their selection probabilities change

To capture developmental changes in movement, we made 2D video recordings of individual larval zebrafish hunting *Paramecia* at 5, 9, and 14 days post-fertilization (dpf) (N=19, 20, and 11 fish respectively) for 15 minutes at 500 fps. The midline of each fish was automatically tracked using a custom image-processing pipeline (Fig. 1a; see Methods), yielding a total of 34,660 swim bouts (92-1367 bouts per fish). Behavior was then classified as either hunting or exploratory via semi-manual annotation (see Methods).

**Figure 1:**
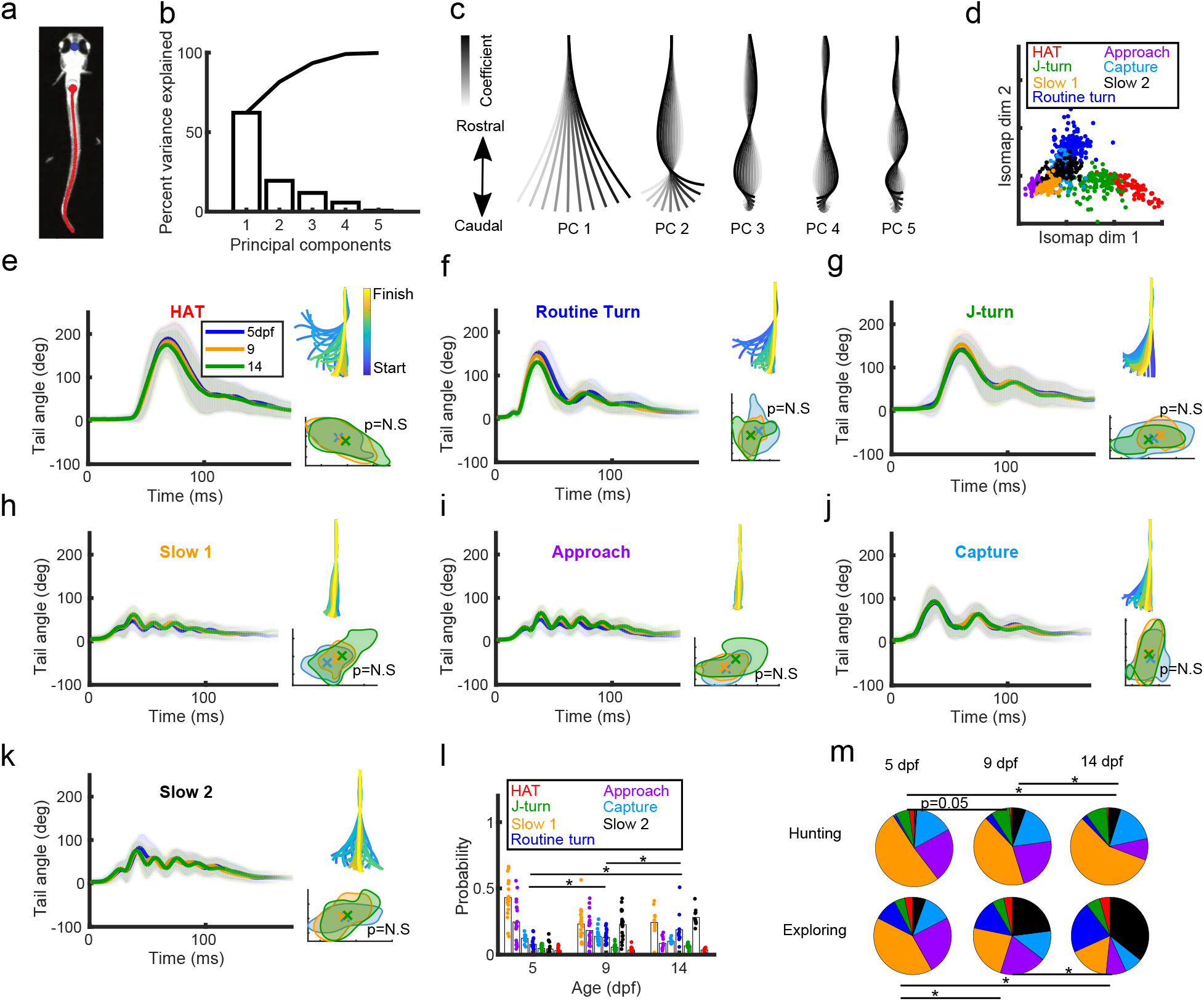
Larval zebrafish bout type shapes are similar over development, however bout type selection probabilities change. **a** In each movie frame the eye midpoint (blue) and 101 points along the tail (red) were tracked. **b** The first 5 PCs explained 99% of the variance in tail curvature. Bars show variance explained by each component, line shows cumulative variance explained. **c** The shapes of the first 5 PCs (eigenfish). **d** Hierarchical clustering on the 20-dimensional Isomap embedding of exemplars. **e-k** Mean and variance of tail-tip angles for each bout type were very similar between 5, 9 and 14 dpf. Thick lines indicate averages across age groups, shading shows 1 std. Top right insets: Example tail traces of bouts for each type (lines drawn every 2 ms). Bottom right insets: For each bout type, there were no statistical differences in the areas covered by age groups in the Isomap embedding space (shuffle test, see Methods). Shaded regions are the outer layers of contour plots for each age group, crosses represent centre of mass for each age group. **l** Fish-averaged bout-type probabilities changed over development (two-sample discrete Kolmogorov-Smirnov test; asterisk indicates P*<*0.0001). **m** Bout-type probabilities changed over development for both hunting and exploratory behavior (two-sample discrete Kolmogorov-Smirnov test; asterisks indicate P*<*0.0001).

To test for developmental changes in movement, we compared several standard kinematic variables. Tail-beat frequency decreased over development (Fig. S1a), while both tail-beat amplitude (Fig. S1b) and tail curvature increased (Fig. S1c). We then sought a representation of movement that would allow finer-scale comparisons. Following the analysis pipeline of Mearns et al [55] we performed Principal Components Analysis (PCA) on local tail curvatures to extract low-dimensional movement features. Pooling all ages, five principal components (PCs) explained *>*99% of tail curvature variance (Fig. 1b), which can be visualized as ‘eigenfish’ (Fig 1c). Performing PCA on each age separately produced no obvious differences in eigenfish or variance explained (Fig. S1d-e). To identify different bout types, Dynamic Time Warping (DTW) [56] was used to compute the similarity between each bout’s trajectory through PCA space (example in Fig. S1f). Affinity Propagation clustering [57] was then used to find representative ‘exemplar’ bouts for the resultant pairwise similarity matrix. Following Mearns *et al.* [55] we then performed a 20-dimensional Isomap embedding on the the exemplar DTW similarity matrix, and used hierarchical clustering on the embedding space to extract 7 clusters. By comparing bout-type shapes with those previously reported [55], we labelled these clusters as High Angle Turns (HATs), J-turns, Routine turns, Slow 1’s, Slow 2’s, Approach, and Capture Strikes (Fig. 1d).

We then asked whether bout movements changed over development. While Slow 1 and Approach bouts changed in both tail-beat frequency and amplitude (Fig. S1g,h), these changes were small relative to the differences between bout types (Fig. S1i,j). Visualizing mean tail-tip angles across time for each bout in a bout type, we found no qualitative differences in structure over development (Fig. 1e-k). Additionally there were no differences in the occupied areas of the Isomap embedding over development (Fig. 1e-k insets). Thus the basic structure of bout types was stable from 5-14 dpf.

An alternative way in which the behavioural space could change is by alterations in bout-type probabilities. Consistent with this, there were significant changes in selection probabilities between 5 and 14 dpf for Slow 1, Slow 2, Routine Turns and Approach bouts, with Slow 2 bouts showing the largest increase and Slow 1 and Approach bouts showing the largest decreases (Fig. S1k). The distributions of bout-type probabilities were different at each age group both when considering hunting and exploring together (Fig. 1l) and separately (Fig. 1m). These changes led to increasing entropy for the distributions of bout-type probabilities, indicating more complexity over development (Fig. S1l). We also examined whether the structure of bout-type transition probabilities changed over development, but found this relatively stable with age compared to the large changes in bout probabilities (Fig. S1m-r). In summary, bout structure remained stable but bout probabilities changed over this period of development.

### A CFD simulation of naturally-swimming fish reproduces velocities and turn angles

Could changing bout-type probabilities over development be related to the changing relationship between the fish and the fluid? To address this we quantified the hydrodynamic interaction between larval zebrafish and fluid using the Immersed Boundary Adaptive Mesh Refinement infrastructure (IBAMR) package [54], which efficiently and accurately models freely-moving fluid-structure interactions. In particular, we adapted the 3D larval zebrafish method used in Bale *et al.* [58].

Larval zebrafish undergo substantial morphological changes between 5 and 14 dpf, including the receding of the fin fold and consumption of the yolk sac [11]. To incorporate these changes into our simulations we volumetrically imaged phalloidin-stained larvae to generate 3D models of an average fish for each age (Fig. S2a-c; see Methods). We also measured and incorporated a developmental increase in fish mass (Fig. S2d). To prepare the experimental tracking data for simulation in IBAMR, we removed all translation in forward/lateral directions and rotation according to the heading angle. During simulations we mapped the relative description of movement for each bout onto the 3D model, similar to Bale *et al.* [58] (see Methods), producing a prediction of the translational and rotational movement for each bout.

We excluded from further analysis bouts that were unlabelled from the clustering pipeline of Fig 1 (n=738) and those whose simulations failed due to numerical instability issues (n=266) (some bouts satisfied both criteria). This left 33,656 bouts in total. We then examined simulation accuracy by comparing CFD-predicted eye-midpoint velocity with experimental eye-midpoint velocity (Fig. 2a; Fig. S2e). First we computed relative error (RE; see Methods) for velocity and excluded bouts where RE*>*0.5 (n=900; Fig. S2f). The remaining 32,756 bouts showed a high correlation between experiment and simulation for mean velocity and turn angle (Fig. 2b,c; Fig. S2g). Simulation accuracy was very similar across age groups (Fig. 2d) and bout types (Fig. 2e). The combined median RE was 0.10. Predicted displacements in the vertical axis were very small, consistent with the experimental recording being in 2D, and were therefore excluded from further analysis (Fig. S2h). Simulation accuracy was insensitive to reducing the number of experimental tracking points (Fig. S2i,j). Examples of the fluid velocity and pressure fields generated by a typical HAT are shown in (Fig. 2f,g), and typical examples of the remaining 6 bout types in Fig. S3. Movies of the CFD velocity field for examples of each of the 7 bout types are provided in supplementary videos 1-7. In summary the IBAMR CFD package accurately reproduces the movement of free-swimming larval zebrafish, and can be used to explore the relationship between larval zebrafish and their fluid environment.

**Figure 2:**
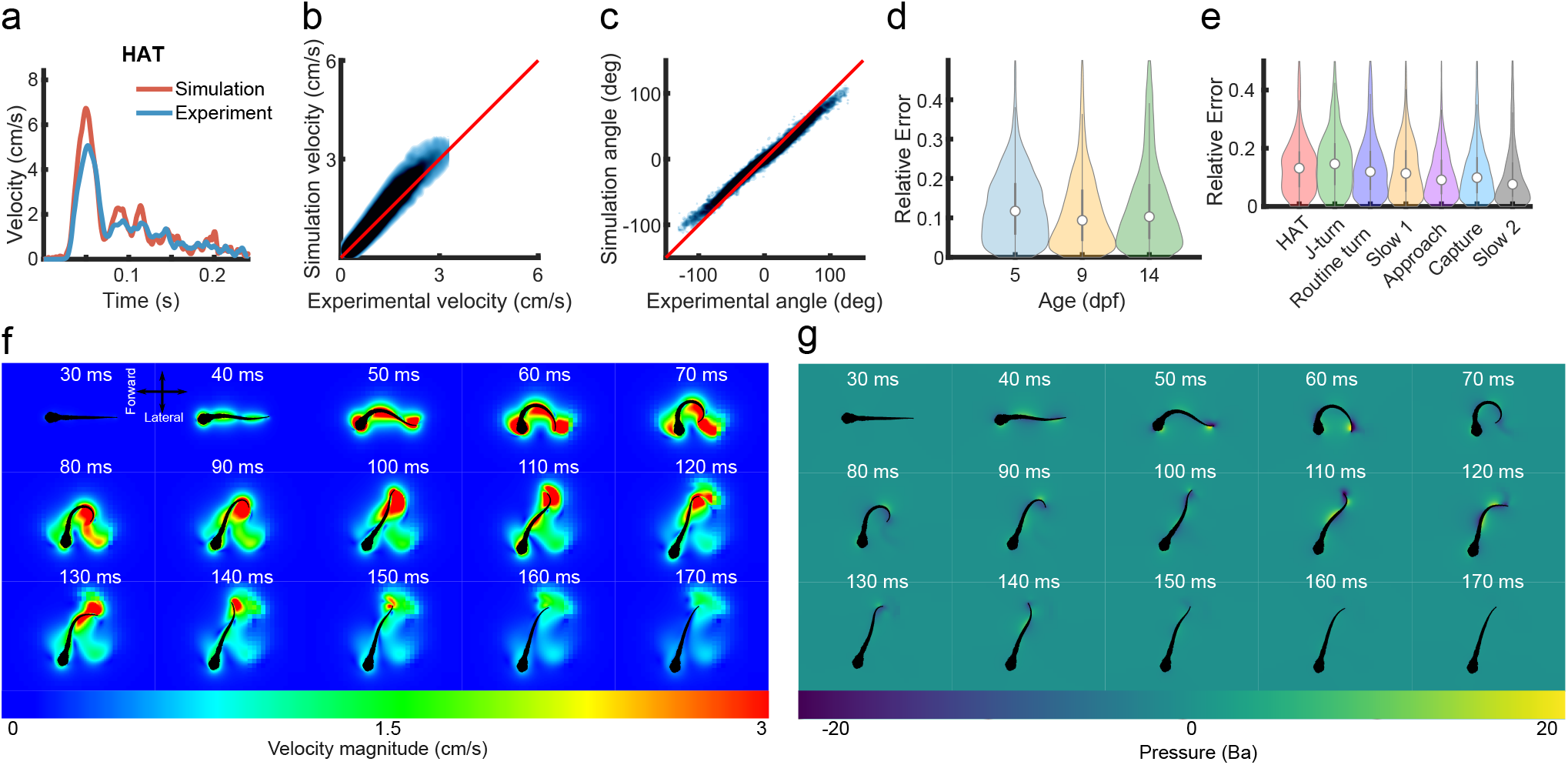
CFD method closely approximates consequences of natural tail movements. **a** Example of the match between simulation and experimental velocities for a 9 dpf HAT (RE = 0.15). **b-c** There was a high correlation between mean experimental and simulation velocities, and between experimental and simulation turn angles, across the entire data set. Blue shaded regions are a contour plot of the experimental values for all bouts plotted against corresponding simulation values, red lines represents a perfect match, R-squared and p-values are from the fit to y = x. **d-e** REs for simulations are stable across both development and bout types. **f-g** Fluid velocity and pressure fields for an example HAT show the tail tip creating vortices in the fluid. T = 0 corresponds to the start of the simulation. Fields are a 2-dimensional z-slice of the 3-dimensional simulation arena. Pressure unit is Baryes (1 Ba = 0.1 Pascals).

### Behavioural space energetics are modulated by fluid regime

How do the energy costs of bouts change over development, and is this related to changing bout-type probabilities? As expected the Reynolds number increased over development (Fig. S4a). However the size of this increase depended on the bout type (Fig. S4b,c), suggesting that development could affect the energy requirements of some bout types more than others.

From each simulation we extracted the hydrodynamic power, which represents the net instantaneous power exerted by the fish into the fluid (Fig. 3a-c). By summing the axis components and integrating over time we obtained the total energy cost per bout. The energies increased over development (Fig. 3d). However again the size of this increase depended on bout type (Fig. 3e,f). Together these results suggest that bout-type efficiencies change differentially over development.

**Figure 3:**
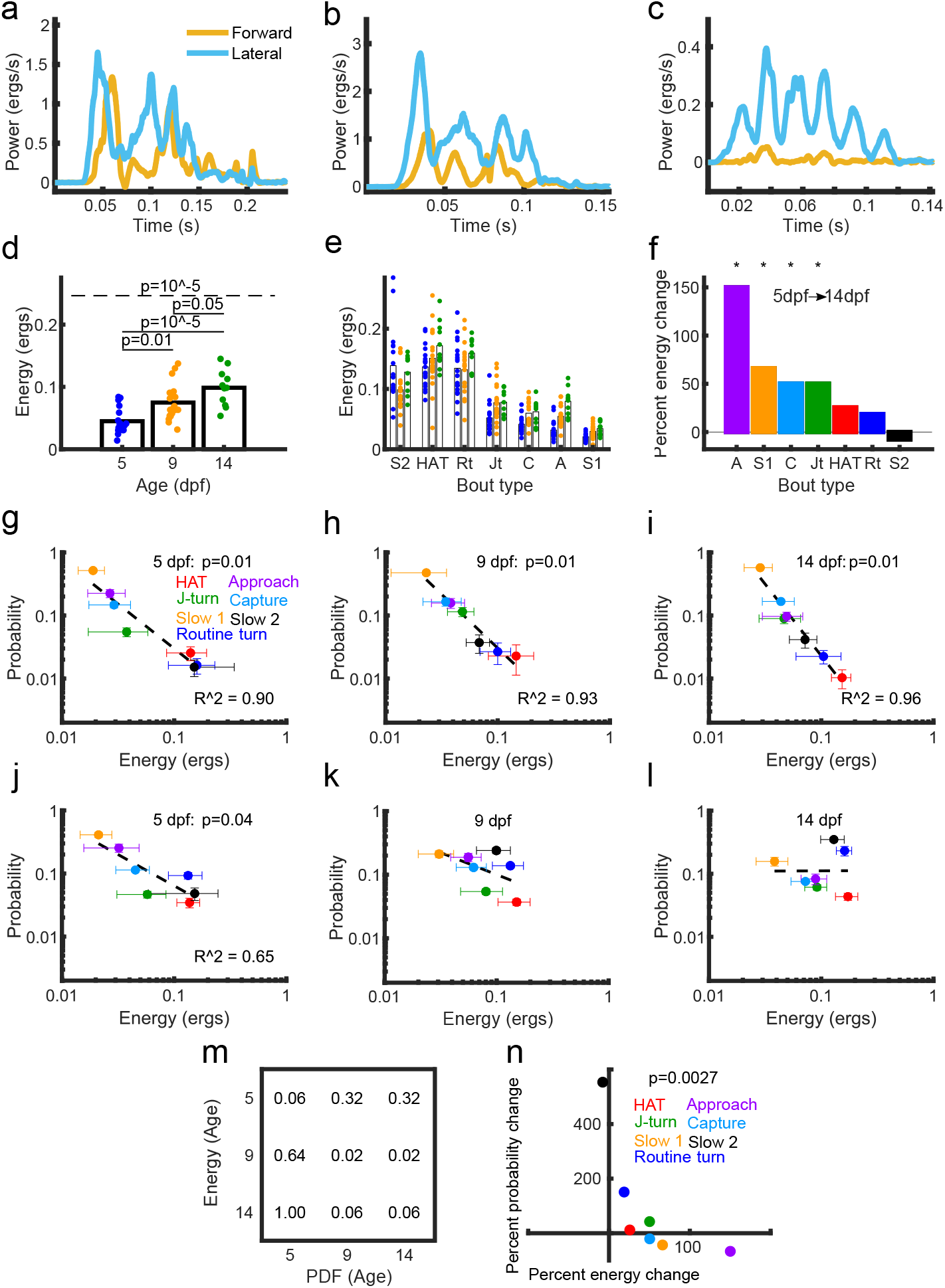
Larval fish maintain a monotonic relationship between selection probability and energy cost despite developmental changes in fluid regime. **a-c** Hydrodynamic power in the forward and lateral directions for an example J-turn (a), Routine turn (b), and Slow 1 (c) bout. **d** Energy cost per bout increases over development. Dashed line gives p-value from ANOVA, solid lines give p-values from pairwise comparisons. **e-f** Energy increases depend on bout type. Dots represent the averages for individual fish. In f, asterisks denote significant difference between 5 and 14 dpf. **g-i** There was a monotonic relationship between bout-type probabilities and bout-type energy costs during hunting behaviour for 5, 9, and 14 dpf. Dots show bout type averages across all fish; p-value is for the Mann-Kendall test for a monotonic trend. **j-l** For exploratory behavior a monotonic relationship was present at 5 dpf but not later stages. **m** A monotonic relationship for hunting was maintained across development via changes in bout-type probability: swapping probabilities and energy costs leads to non-monotonic relationships (Bonferroni-correction n=9, hence diagonal entries are higher than in g-i). **n** There was a monotonic relationship between percentage energy and probability change, such that bout types with the largest energy increases had the largest decreases in probability between 5 and 14 dpf.

We then asked how energy constrains behavior. We first separated exploratory (Fig. S4d) and hunting behavior (Fig. S4e), and found a developmental increase in energy cost for exploratory but not hunting behavior. There was an increase in Capture cost (mean energy cost for capture events, Fig. S4f), but no changes in Total cost (sum of energies for all hunting events divided by number of *Paramecia* captured, Fig. S4g), Event cost (mean energy cost for hunting events, Fig. S4h) or Abort cost (mean energy cost for abort/miss events, Fig. S4i). This was despite an increase in fish mass and in the energy required for a J-turn to produce a particular angular change (Fig. S4j).

Remarkably, during hunting, there was at all ages a monotonic, power-law relationship between bout-type probability and bout-type energy cost, such that more expensive bouts were used less often (Fig. 3g-i). This relationship was robust to manipulations of the number of clustered bout types (n = 4-10 bout types; Fig. S5a), and was also present when energies were binned rather than averaged across bout types (Fig. S5b). The monotonic relationship during hunting was also present for the majority of fish considered individually (Fig. S6). In contrast for exploratory behavior, while there was a monotonic relationship at 5 dpf, this was not present at later developmental stages (Fig. 3j-l). This non-monotonicity was robust to different numbers of bout types (Fig. S5c) and binning methods (Fig. S5d), but the monotonicity at 5 dpf was only present for 7 bout types (Fig. S5c).

Over development both probabilities and energy costs changed. To determine whether these behavioral changes facilitated conservation of the monotonic relationship during hunting, we randomly mixed bout-type costs and probabilities between age groups. The monotonic relationship was not present when mixing bout-type energies at 14 dpf with bout-type probabilities at 5 dpf (Fig. 3m). Together this suggests that bout-type utilization depends on energy costs. Consistent with this hypothesis, there was a negative correlation between percentage increase in energy cost and percentage increase in bout-type probability (Fig. 3n). Together these results suggest that larval zebrafish hunting adapts to changing energy constraints over development.

### Approximating CFD energy

So far all energies quoted were based on CFD simulations, which require non-trivial effort. We now asked to what degree simpler calculations could approximate CFD energy. We began by testing basic kinematic features, and found that tail frequency and tail amplitude had very low correlations with CFD energy usage (Fig. S7a-b). Tail vigor [59], defined in terms of the standard deviation of the tail angle, was also a poor predictor (Fig. S7c,d). We then tested the ”total kinetic energy” (TKE), defined as *TKE* = <u>^1^</u> *mv*^2^ where *m* is fish mass and *v* the average translational velocity of the fish’s eye midpoint during the bout. TKE had a medium correlation (*R*^2^ = 0.68) with the CFD energy usage (Fig. S7e), with a scaling term of 202. We then wondered if the approximation could be improved with a machine-learning approach. We trained a Gaussian Process Regression model (GPR) to predict CFD energy usage for a swim bout based on the absolute eye-midpoint displacement for that swim bout, using Matlab’s *fitrgp* function. Training on 70% of the dataset (all ages simultaneously) and then testing on the remaining 30% produced a high correlation with CFD energy (*R*^2^ = 0.80, Fig. S7f), with a median relative error compared to the CFD values of 0.205. A more accurate model was achieved by using the final x and y displacements of the eye midpoint as training features for each swim bout (*R*^2^ = 0.85, Fig. 7g), with a median relative error of 0.196. However the ratio of CFD energy to TKE differed with age and with bout type (Fig. S7h). In particular bouts producing primarily forward motion (Approach, Capture, Slow 1 and Slow 2) required more CFD energy compared to fish kinetic energy than turning bouts (J-turn, Routine turn, HAT). Thus, although being correlated at the population level, approximations of CFD energy based only on displacements may be unreliable at the single-bout level.

### Larval zebrafish reared in increased-viscosity water have altered behavioural spaces

The behavioral changes we observed over development could arise from either pre-specified developmental programs, or through active adaptation to the changing fluid environment as the fish grows. To test whether larval zebrafish can adapt to novel conditions, we reared larval zebrafish (n = 30) from 1 dpf in water where the viscosity had been increased by a factor of 3.3 (see Methods) by adding a small amount of dextran (“viscous water”), and compared hunting at 8 dpf in viscous water (n = 15 fish; n = 12,970 bouts; 448-1116 bouts per fish) versus hunting at 8 dpf in normal water (“water”; n = 15; n = 16,095 bouts; 654-1463 bouts per fish) (Fig. 4a). We compared these results with the 9 dpf data collected for Fig 1, where the fish were both reared and tested in water.

**Figure 4:**
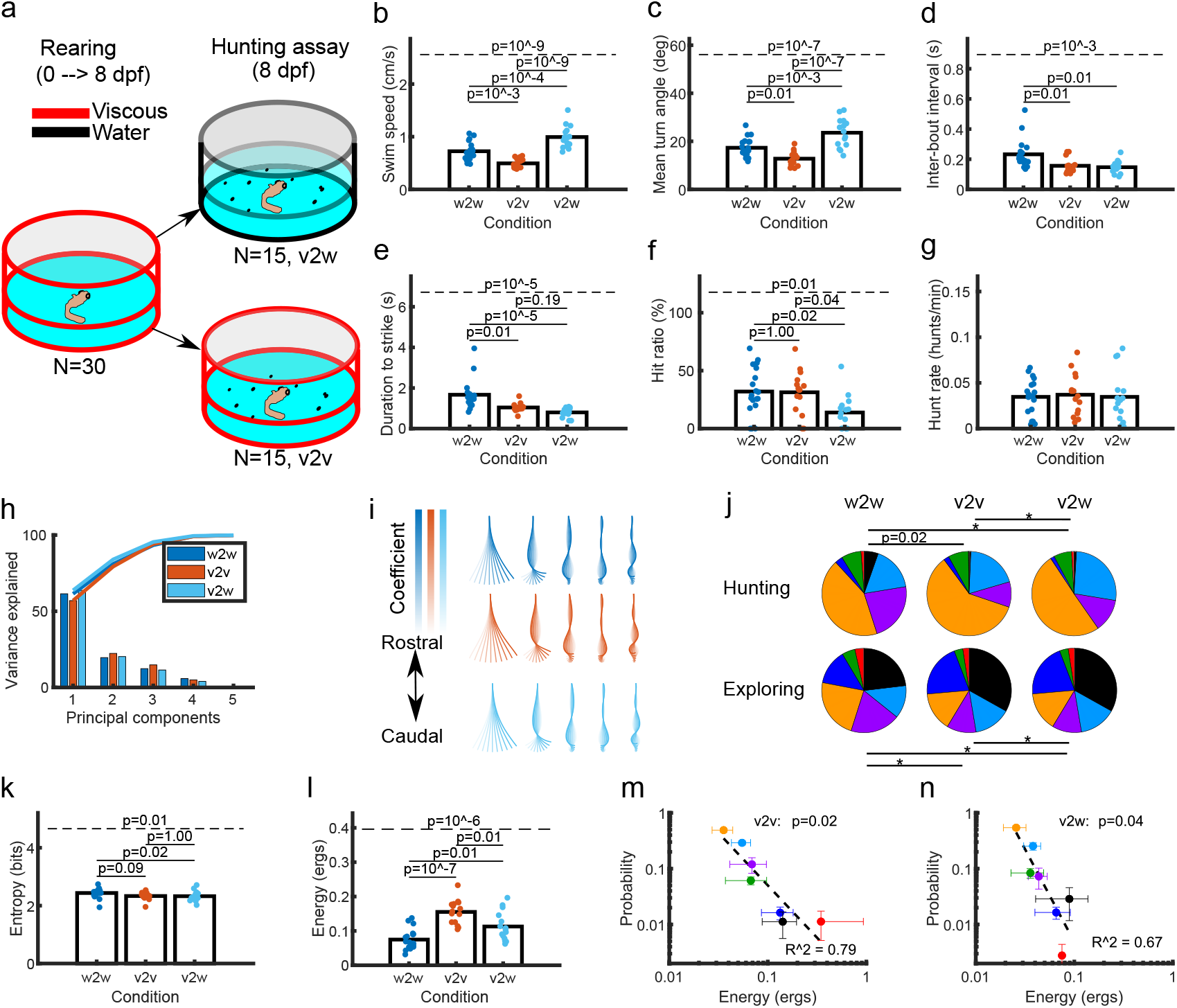
Viscous-reared larval zebrafish have altered behavioral spaces. **a** Fish were initially reared in viscous water from birth, transferred to either water (v2w) or viscous water (v2v) at 8 dpf, and then recorded hunting *Paramecia*. **b-d** The v2v fish had lower speeds, turning angles, and inter-bout intervals than w2w fish. Dashed lines indicate ANOVA test across conditions, solid lines indicate pairwise comparisons. **e-g** The v2v fish had lower hunt durations but similar hit ratios and hunt rates compared to w2w fish. **h** The variance explained by the leading principal components was similar across conditions. **i** The shapes and orderings of the eigenfish were similar across conditions. **j** There were differences in the distributions of bout-type probabilities between w2w and v2v fish, for both hunting and exploring behavior (two-sample discrete Kolmogorov-Smirnov test; asterisks indicate P*<*0.0001). **k** The behavioral complexity was smaller for v2w fish than w2w fish. **l** Both the v2v and v2w fish used more energy per movement than w2w fish. **m-n** There was a monotonic relationship between bout-type energy and probability for both v2v and v2w fish respectively (monotonicity tested via Mann-Kendall). Dashed lines are the fitted linear regression *y* = *β*_0_ + *β*_1_*x*.

Consistent with viscous environments being more challenging for movement, viscous-reared*→*viscous-tested fish (v2v) had smaller swim speeds, turn angles, and shorter interbout intervals than water-reared*→*water-tested fish (w2w; Fig. 4b-d). The v2v fish had shorter hunting event durations (Fig. 4e) yet similar hit ratios and hunt rates (Fig. 4f,g) to w2w fish. The v2v fish also hunted *Paramecia* that were closer and slower moving than w2w fish, from the same detection angle (Fig. S8a-c), leading to shorter strike distances and a non-significant difference in the number of bouts per hunting event (Fig. S8d-e). The v2v fish also had larger undershoots than w2w fish (Fig. S8f), defined as the change in angle between fish heading direction and the prey prior to the first hunting movement.

Consistent with the hypothesis that fish are calibrated to their environments, and are thus miscalibrated when first placed in a new environment, viscous-reared*→*water-tested fish (v2w) had higher swim speeds, turn angles, and lower inter-bout intervals than w2w fish (Fig. 4b-d). Furthermore, compared to w2w fish, v2w fish had shorter duration to strike (Fig. 4e), lower entropy (Fig. 4k), were less successful at catching prey (Fig. 4f), initiated hunting at shorter distances (Fig. S8a), and had shorter distances (Fig. S8d) and fewer bouts (Fig. S8e) to strike. Together this suggests the possibility of predictive coding/internal models of the environment that depend on viscosity.

Given these changes in hunting strategy, we investigated whether they were related to changes in movement mechanics or selection probabilities. First we generated separate behavioural spaces for each rearing condition, and found no obvious differences in variance explained (Fig. 4h) or between eigenfish (Fig. 4i). We then generated a combined behavioral space and found no difference in the distribution of conditions within the Isomap embedding (Fig. S9a-b). In the combined PC space, we then trained a support vector machine (SVM) model to label v2v and v2w bout types given the pre-labelled w2w bout types (see Methods). When comparing each bout type across conditions, there were few differences between mean tail-tip traces (Fig. S9c). v2w fish had higher frequencies than w2w fish (Fig. S8g-i), while tail-beat amplitude did not change between conditions (Fig. S8j-l). Therefore, viscous-reared fish had similar movement mechanics to water-reared fish.

In contrast, there were differences in bout-type probabilities between conditions during both hunting and exploring (Fig. 4j), with viscous-reared fish (i.e. v2v and v2w) using fewer J-turns and more Routine turns during exploration, and using more Capture Strikes during hunting, than water-reared fish (Fig. S8m-o). v2w fish had lower behavioral complexity than w2w fish (Fig. 4k). Overall these results show that larval zebrafish adapt behavioral strategies such as bout-type probability in response to altered environmental constraints.

We then asked whether these changes were related to energetic considerations. Consistent with viscous water being more challenging for movement, v2v fish (calculated from the CFD simulations using appropriately increased viscosity) had higher energy usage than w2w fish (Fig. 4l). v2w fish also had higher energy usage than w2w fish, presumably due to miscalibration with the environment. There was a non-uniform increase in energy usage for each bout type (Fig S8p), with HATs and Routine turns being the most affected and Approach bouts the least. Compared to the w2w case capture costs were higher but abort costs were the same for the v2v case (Fig S8q,r). However, remarkably, the monotonic relationship between bout selection probability and energy cost during hunting was maintained in both v2v and v2w fish (Fig. 4m-n). Overall these results suggest that viscous-rearing adaptations act to limit increases in hunting costs.

### Larval fish adapt to altered-viscosity fluid within 24 h

Above we examined fish behavior immediately after long-term rearing in viscous water. We now asked whether viscous-reared fish could adapt to water within 24 h. We reared a new cohort of fish in viscous water and then, as a baseine measurement, recorded hunting at 8 dpf in water for 15 min (N=24 fish, N=26,673 bouts, 189-2221 bouts per fish; see Methods); we refer to these results as “8v2w”. We then kept half of the cohort in water for 24 h, while the other half were kept in viscous water for 24 h. After this, both groups were recorded hunting in water (Fig. 5a); we refer to these as “9w2w” (N=11,828 bouts, 340-1490 bouts per fish) and “9v2w” (N=12,655 bouts, 412-1739 bouts per fish) respectively.

**Figure 5:**
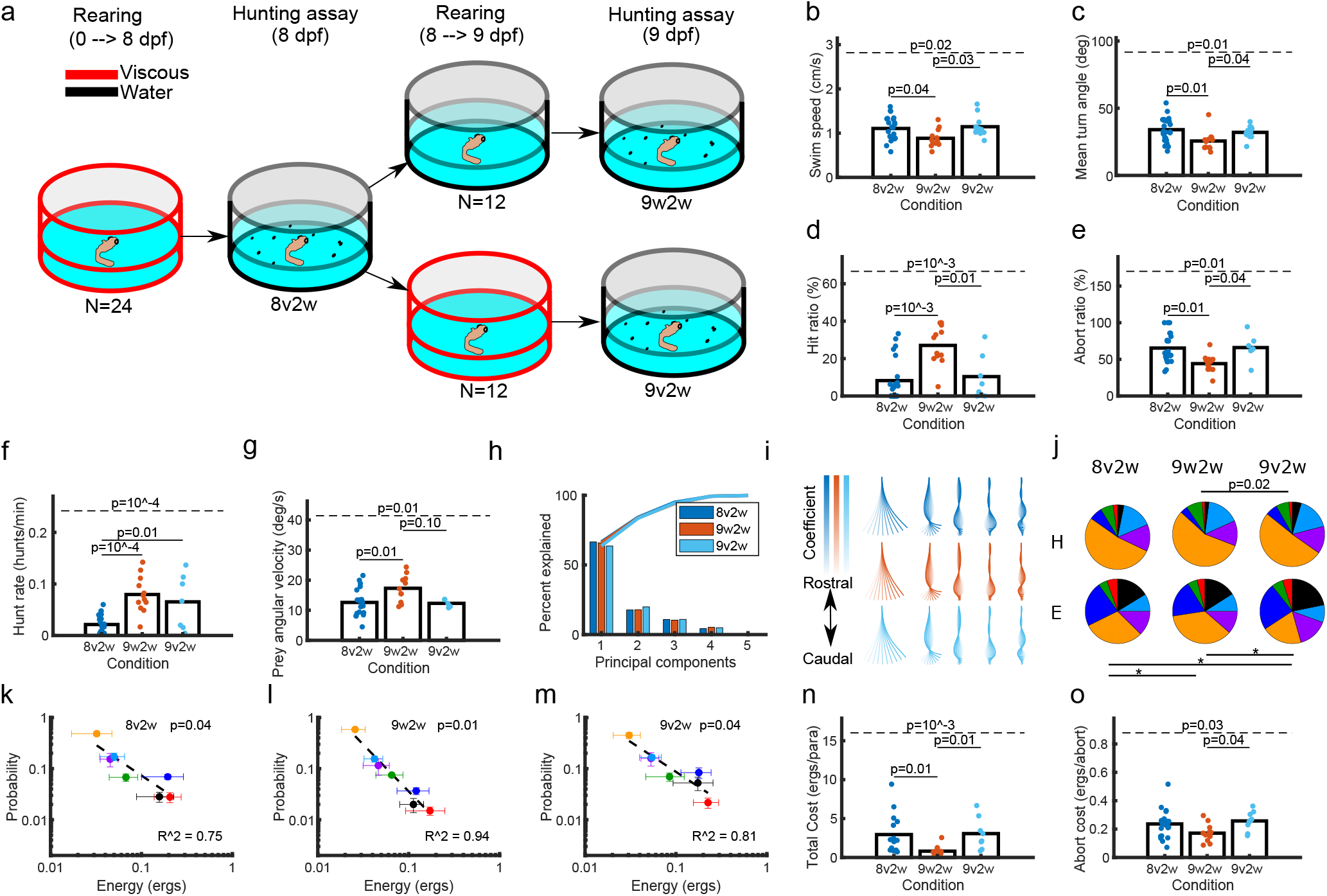
Larval fish adapt to altered-viscosity fluid within 24 h. **a** Experimental design. **b-c** 9w2w fish swim speeds and turning angles were smaller than 8v2w and 9v2w fish. **d-g** Hit ratio, abort ratio, hunt rate, and prey angular velocity were different between 9w2w and 8v2w/9v2w. **h** The variance explained by the first five principal components was similar between conditions. **i** The eigenfish had similar shapes and orderings across conditions. **j** There were differences in the distributions of bout-type probabilities between 9v2w and 9w2w fish, for both hunting and exploring behavior. **k-m** There was a monotonic relationship between energy and selection probability in each case. **n** 9w2w fish expended less energy per paramecium during hunting events compared to 8v2w and 9v2w fish. **o** There were differences in the energy per abort event between 9w2w and 9v2w fish.

The 9w2w fish had lower swim speeds and turn angles than 8v2w and 9v2w fish (Fig. 5b-c). The hunting metrics of 9w2w were also different from 8v2w and 9v2w fish, with 9w2w fish having higher hit ratios, lower abort ratios and higher hunting event rates, and also selecting prey with higher angular velocities (Fig. 5d-g). There were no differences in prey detection angle or the number of hunting bouts between conditions (Fig. S10a-b). Thus hunting adaptation occurred within 24 hours of exposure to an altered fluid environment.

We then asked whether movement mechanics and/or bout-type probabilities were altered. When the behavioural spaces for 8v2w, 9w2w and 9v2w were generated separately there were no obvious differences in the variance explained (Fig. 5h) and no qualitative differences between eigenfish (Fig. 5i). We then generated a combined behavioral space and found no difference in the distribution of conditions within the Isomap embedding (Fig. S10c). We used a SVM model as described above to label the bout types and found few differences in tail-tip traces between conditions (Fig. S10d). In particular there were no differences in either tail frequency or amplitude for all bout types when comparing 8v2w and 9w2w (Fig. S10e,f). However there were significant changes in bout-type probabilities between conditions during both hunting and exploring (Fig. 5j, Fig. S10g). Thus, behavioral adaptation via altered selection probabilities occurred within 24 h.

We then wondered whether these behavioral changes were related to energetic considerations. In each condition the monotonic relationship between energy usage and probability for bout types was maintained (Fig. 5k-m). However the 9w2w fish had much lower Total Cost (energy per *Paramecia*) than 8v2w and 9v2w fish (Fig. 5n) and lower abort costs (Fig. 5o). This suggests that fish adapt behavioural strategies to their environment in ways that reduce energy consumption, and that these adaptations can occur on timescales as short at 24 hours.

## Discussion

Here we combined detailed behavioral analysis of larval zebrafish with CFD simulations to uncover how the environment constrains behavior during early development. Behavioral adaptation occurred primarily through changes in bout-type probabilities rather than movement mechanics, as demonstrated through both developmental and environmental perturbations. There was a robust monotonic relationship between energy cost and selection probability for each bout type, that was maintained regardless of condition. These adaptations served to reduce energy costs, particularly in the case of hunting behavior, and occurred within 24 hours of exposure to novel conditions.

### Bout types over development

Previous studies have shown major changes in larval zebrafish movement mechanics between 3 and 5 dpf. At 3 dpf zebrafish use cyclic swimming, where the tail constantly beats, but then transition into burst-and-coast swimming by 5 dpf, where there are beat and glide phases [45]. Here we showed that over the subsequent 10 days of development, the bout structures comprising burst-and-coast swimming remained stable while bout-type probabilities changed. Comparing across species, Dunn *et al.* [60] found that between 7 and 30 postnatal days rats added body-grooming and rearing behaviors to their behavioral repertoire [60]. Dunn *et al.* [60] also showed an increase in behavioral complexity over development, which is similar to our findings. In contrast the behavioral complexity of *Drosophila* is largely unaltered between 0 and 70 days of age [61], while the behavioral complexity of humans has an inverted U-shape trend over development [62]. Comparing to other zebrafish studies, it is known that adult zebrafish display more movement types than typically identified at larval stages, and from 14 dpf onwards zebrafish bouts become more connected so that by 30 dpf there is complete fusion of most hunting events [63]. However comparisons between these studies can be challenging, since fish size and age are not necessarily perfect proxies for zebrafish development [64]. Ultimately, behavioral complexity across organisms depends on both task complexity and the number of unique tasks an organism performs.

### Large-scale CFD simulations of naturalistic movements

Using the IBAMR package, we performed the first high-throughput 3D CFD simulations of the larval zebrafish behavioral space. Bale *et al.* [58] previously used IBAMR to simulate larval zebrafish with experimentally-derived movement, however they transformed the original burst-coast motion into steady/cyclical movement before simulation, and only analysed one forward swim bout. In comparison we preserved burst-coast swimming and analysed *>*100,000 turning and forward swim bouts. Despite these extensions, our CFD simulations achieved a median RE 0.10, compared to 0.18 in Bale *et al.* [58]. Our power expenditures were in the same range as Bale *et al.* [58], who computed 1.67 erg/s for steady swimming at approximately 1.2 cm/s, though ours are generally slightly lower, likely due to steady movement requiring more power. Zhao *et al.* [65] developed a CFD method to achieve high-throughput simulations of larval zebrafish, however the analysis was restricted to quasi-steady high-speed cruising movements and excluded sudden starts. Li *et al.* [66] developed a CFD method to simulate sudden-start escape bouts in larval zebrafish, however the full behavioral repertoire was not considered.

Our simulations did not consider the pectoral fins. It has been argued that these fins do not play a significant locomotor role in larval zebrafish, but instead have a primarily respiratory function [67]. Evidence for this includes that the swimming kinematics of larval zebrafish lacking pectoral fins as a result of genetic mutation are similar to those of normal larvae [68]. Furthermore fluid dynamics simulations show that fin movements are well-suited for posterior transport of fluid along the body surface [69]. More recently it has been shown that the pectoral fins are important for vertical motion [70], however we did not image 3D motion in our assays. Our simulations also ignored elastic, dampening, and muscle-activation forces within the fish tail. Tytell *et al.* [71] created integrated neuromechanical models of the lamprey where these forces combined to drive movement, however movements were limited to cyclic forward motions. A future challenge is to reproduce naturalistic motion of zebrafish using biologically-realistic mechanics.

### Connection between energy change and behavioral change

We demonstrated that different bout types have different energy costs, and these costs change differentially between 5-14 dpf. We also showed that these changing energetics were reflected in the selection probabilities of different movements, such that there was a negative correlation between the increase in bout-type energy cost and increase in bout-type probability. This extends previous work investigating the energetics of burst-and-coast versus cyclical swimming between 1-5 dpf, where it was theoretically predicted [72] and experimentally demonstrated [14, 48] that fluid-regime alterations of drag profiles for differing swim styles could explain the change from continuous to burst-and-coast swimming. Our study shows that this adaptation mechanism is not limited to early development, and may be maintained until at least juvenile stages. The finding that behavioral changes match movement-efficiency changes has also been found for adult *Drosophila* [61], suggesting an underlying principle across species that minimizing energy costs plays an important role in behavioral changes.

Optimal search theory predicts a Levy-Flight strategy when prey is sparse and randomly distributed [73]. This has been shown theoretically to maximize prey encounter rates [74] and has been observed in several animals [73, 75]. Our work adds to this literature by considering for the first time experimentally-measured energy costs rather than simply displacements. Our study shows that energy itself may be Levy-Flight distributed. However in our assay *Paramecia* are relatively dense (initially roughly 4-5 *Paramecia* in the fish’s field of view at each moment), are constantly moving, and their density decreases over time, which does not match the assumptions underlying the optimality of Levy Flights.

### Mechanical efficiency vs metabolic efficiency

Li *et al.* [51] found that larval zebrafish minimize COT during rheotaxis by altering tail frequency and amplitude, using a mechanical approximation for power. Similarly, Mandralis *et al.* [48] analysed the efficiency of escape manoeuvres under given energy budgets using mechanical approximations. Our study extends these works by analysing the efficiency of larval zebrafish hunting, a multi-movement behavior. However, mechanical efficiency and metabolic efficiency are not necessarily the same. It has been predicted theoretically that there is an U-shaped relationship between mechanical energy and metabolic energy such that low and high cruising speeds require similar metabolic energy costs [76]. However until there is more empircal support for this, mechanical energy remains a reasonable first approximation.

### How do larval zebrafish encode energy costs?

An important question for future work is how fish monitor energy usage. While fish lack the proprioceptive structures like muscle spindles and Golgi tendon organs found in mammals, they do have sensory systems that perform similar functions. Mechanosensory feedback via spinal microcircuits contributes to controlling locomotor frequency [77], and a central organ of proprioception in the zebrafish spinal cord was recently discovered that senses spinal bending and helps control locomotor rhythm [78]. However this organ only develops after the time period we have studied here. Results from virtual reality arenas demonstrate that altering visual feedback in fictive swimming can cause adjustments of swim strength [79]. However these changes can occur in paralysed fish, and so do not rely on direct measurements of energy consumption. Water-flow sensing in fish occurs via the lateral line system [80], which can subserve hydrodynamic imaging of the environment [81]. Cranial neuromasts (part of the lateral line system) have been shown to be important for prey hunting in the dark [82], but only in fish older than those we have studied here. However it is unclear how measurements of water flow by themselves would be sufficient for estimating energy consumption. Metabolic processes such as the rate of oxygen consumption and the accumulation of metabolic byproducts can serve as indicators of energy cost [83], but whether these underlie the adaptation of bout probabilities that we have observed remains to be determined.

### Energetic neuroadaptation

Together our results suggest a new field of study: how neural circuits encode and adapt to changing energy costs of movement in order to optimize behavior. Fish offer an attractive entry point for this, due to the relative ease of calculating the mechanical energy expended in fluid environments. However future work in rodents and other legged species may uncover relevant approximations for energy, opening up a broad range of species for the investigation of how energy constrains behavior.

## Materials and Methods

### Zebrafish

All experimental procedures were performed with approval from either The University of Queensland Animal Ethics Committee, or the Animal Studies Committee at Washington University in St Louis under IACUC protocol 21-0143, and adhere to NIH guidelines. Fish embryos were raised in E3 embryo medium at 28.5°C on a 14/10 h light/dark cycle. From 1 dpf larvae were kept in small groups in 100 mm Petri dishes, and were fed live rotifers (*Brachionus plicatilis*) daily from 5 dpf.

### Hunting-behavior assay

Individual fish were placed into a feeding chamber (diameter: 20mm; depth: 2.5mm; CoverWell Imaging Chambers, catalogue number 635031, Grace Biolabs, USA) filled with E3 medium with depth of 2.5mm and 30-35 *Paramecia* (*Paramecium caudatum*). The chamber was placed onto a custom-made imaging stage consisting of a clear-bottom heating plate at 29.5°C, an infrared LED ring below (850 nm, LDR2-100IR2-850-LA powered by PD3-3024-3-PI, CCS Inc., Japan), and a white LED ring above (LDR2-100SW2-LA, CCS). Images were recorded using a CMOS camera (EoSens 4CXP, Mikrotron, Germany) at 500 fps using StreamPix (NorPix, Canada). Recording of hunting behaviour started after the first attempt for feeding was made by the fish, and each fish was then recorded for 10-15 mins.

### Increased-viscosity assays

For viscous rearing, fish embryos were raised as described above except for the addition of 3% dextran. To measure fluid viscosity, we used a microviscometry method established by Bishop *et al.* [84].

### Analysis of feeding events

Automated tracking of the fish and *Paramecia* was performed using custom image processing software in MATLAB as detailed in Avitan *et al.* [17], with minor modifications. In brief, frames were first pre-processed to remove the static background using a Gaussian background model. The approximate location of the fish was identified by connected-components analysis on the resulting foreground mask. The position and orientation of the fish were calculated by tracking the midpoint between the eyes and the centre of the swim bladder. This was achieved using a set of correlation filters on pixel values and histogram of oriented gradients features. Filters were rotated through 0,5,10,…,360 degrees and scaled through 60,65,70,…,100% with respect to maximum fish length, to accommodate changes in heading angle and pitch respectively. Filters were trained by manual annotation of the two tracking points in ten randomly selected frames for each fish. Bout timings and tail kinematics were calculated by first performing morphological thinning and third-order Savitsky-Golay smoothing to extract 101 evenly spaced points along the midline of the tail. Individual bouts were segmented by applying a manually-selected threshold to the amplitude envelope of the mean angular velocity of the most caudal 20% of tail points. Prior to applying the threshold, the angular velocity time series was smoothed using a low-pass filter. The amplitude envelope was estimated using a Hilbert transformation.

Detection of *Paramecia* was performed using connected components analysis to extract the location of prey-like blobs in each frame from the foreground mask. Multi-object tracking of *Paramecia* between frames was achieved using Kalman filtering and track assignment, which enabled tracking through collisions and short periods of occlusion.

Hunting events were defined as eye convergence followed by at least two bouts before deconvergence. The start frames of eye convergence were detected manually for experiments of Figs 1 and 3, and were detected automatically by a pre-trained classifier for experiments of Figs 4 and 5. An event classification pipeline was developed using a Convolutional Neural Network (CNN) to identify frames where the eyes were converged. A dataset of 132 manually-annotated feeding assays was used for training. To prepare the data for classification, the fish head was extracted from each frame using the tracking pipeline described above. The resulting images were then cropped to dimensions of 64×64 pixels, rotated to the same direction and converted to grayscale. Random adjustments to image brightness were applied to augment the training data. The dataset was divided into 70% for training, 20% for validation, and 10% for testing. For the training and validation sets, the middle frame of hunting events was designated as class 1 (converged frame), while random frames without hunting events were labeled as class 0 (deconverged frame). This ensured a balanced representation of both classes in the training and validation datasets.

The CNN classifier was based on the efficientnet-b0 architecture [85]. Modifications were made to the input layer to adapt the network to the 64×64 input image size. The model was initialized using pretrained weights obtained from the ImageNet dataset. During the training phase, a batch size of 32 was used, and the model was trained for 10 epochs. To prioritize the detection of converged frames (class 1), weight of 0.9 were assigned to converged frames, and 0.1 to deconverged frames. The model’s performance was evaluated based on its accuracy on the validation set, and the best-performing model, which achieved 93% accuracy, was selected as the final model. The final model was evaluated on the test set, comprising 14 feeding assays. In contrast to the training and validation sets, which utilized subsampled frames, evaluation on the test set included all frames within each assay to assess the classifier’s ability to detect eye convergence events. The model achieved an average false negative rate of 2% on the hunting events in the test set.

Events were then manually classified based on whether the fish aborted pursuit of the target paramecium (abort event), pursued but failed to capture the target (miss event), or the fish successfully captured the target (hit events or spit events). Hit ratio was defined as the number of hits divided by the number of hunting events, and abort ratio as the number of aborts divided by the number of hunting events. Event end was determined by eye deconvergence for abort events, and for other events by the end of the strike bout. The target paramecium was defined as the nearest paramecium towards which the first tuning bout was made.

### Generating the behavioural space

For water-reared experiments, we first excluded any bouts or hunting events where the fish came within 0.25 mm of the dish boundary. To exclude tail-tracking errors when the fish rolled, encountered debris, or had *Paramecia* collisions, we removed bouts with displacement less than 5 pixels (*≈*0.1 mm), bouts for which the tail length differed by more than 2 standard deviations from the mean during more than 35% of the bout, and bouts where angular differences larger than 30 degrees between eye and swim bladder comprised more than 5% of the bout. Remaining bouts containing proportionally fewer tail-length or heading-angle errors were smoothed via linear interpolation. In total 11,880 bouts were removed by these filters, leaving n=34,660 bouts in total (n=10,350 for 5 dpf, n=17,517 for 9 dpf, and 6793 for 14 dpf). For viscous-reared experiments, we used similar exclusion criteria. In Figure 4 there were 12,548 bouts removed by these filters leaving n=29,065 bouts in total (n=16,095 for v2w fish, and n=12,970 for v2v fish). For Figure 5 there were 23,613 bouts removed by these filters leaving n=51,156 bouts in total (n=26,673 for 8 dpf v2w fish, n=11,828 for 9dpf w2w fish, and n=12,655 for 9 dpf v2w fish).

The behavioural-space analysis pipeline was adapted from [55]. To transform bouts into postural dynamics, PCA was performed on tail curvatures from the swim bladder to the tail tip. Data were normalised by subtracting the mean tail curvature and dividing by the standard deviation. PCA was first performed on each age-group to create separate PC spaces, and then performed on all age-groups together to create a combined PC space.

To classify bout type, we first calculated the distance between each bout’s PC trajectory via dynamic time warping (DTW), using a warping window of 10 ms and the first five PCs in the combined PC space. For bouts of different lengths, the shorter-length bout was padded with zeros to match the length of the larger bout. Left/right polarity was removed by performing each alignment twice, with all values reversed for one trajectory the second time, and taking the minimum of the pair as the similarity. Affinity propagation was then performed on the similarity matrix, with the median similarity between bouts used as the preference for the clustering. Initially 1733 exemplars were identified; we then excluded any exemplar containing less than 3 bouts, leaving 1111 exemplars. Isomap embedding into a 20-D space ([86]) was then performed on the remaining exemplars, by using the DTW distances to construct a nearest-neighbors graph (nearest neighbors = 20), calculating minimum distances between pairs of points, and finding the eigenvectors of the graph distance matrix. Hierarchical clustering was used to classify bout types.

For viscous-reared fish we created a combined PC space from viscous and water-reared data. For Figure 4, we used the Figure 1&4 fish; for Figure 5, we used the Figure 1&5 fish. We then used the prior-classified water-reared fish to train a Support Vector Machine model to identify bout types from PC trajectories within the combined PC space. With the trained model, we then predicted bout types of the viscous-reared PC trajectories within the combined PC space. The SVM was trained on 5-D PC trajectories (unrolled, each component zero-padded to the length of longest trajectory) and bout-type labels, using a random 80/20 training/test split. Viscous-reared bouts with longer trajectories than the longest water-reared bout were ignored. We used the sklearn.svm function SVC, with C=1 and balanced weight class. The test set had an adjusted rand index of 0.61.

To compare distributions of rear/test conditions within a combined Isomap embedding, we used a shuffle test based on overlapping areas. First we made KDE plots for each rear/test condition in the first two dimensions of Isomap space, then selected the outermost contour for each condition and computed the overlapping area. This represented experimental similarity within the combined Isomap space. To represent chance levels of similarity, we shuffled rear/test conditions (10,000 shuffles) and recomputed the overlapping areas. A significant difference in Isomap embedding distribution occurred when the experimental overlapping area was smaller than the 5th percentile of the shuffled overlapping areas.

### CFD preprocessing

To address CFD stability issues, the recorded frame rate of 500 fps was increased to 24,500 fps via linear interpolation of 50 points between each time step. To smooth small-scale tracking noise we applied a Kalman filter to the interpolated data. To extend the tracking points to the full length of the larval zebrafish, the head regions were assumed to be rigid and linear interpolation was applied between the midpoint of the eyes and the swim bladder using similar spacing to the original tracking. This resulted in N=125 evenly spaced points describing the fish’s mid-line.

Since the CFD method used relative motion to predict global motion, we removed net forward, lateral, and rotational movement for the duration of each bout. Net rotation was defined by the heading angle of the fish, between the eye midpoint and the 25th point along the midline. Net forward and lateral movement were defined via the initial heading angle of the fish and the eye-midpoint position. We saved the displacement/rotation information for validation during post-processing. Then, local tail curvature was calculated between each midline point for consecutive time steps.

Each swim bout was simulated separately, and each simulation took approximately 1.5-3 hours on an Intel Cascade Lake core housed in Washington University’s Research Infrastructure Services Scientific Computing Platform (usually we ran batches of several hundred simulations in parallel). The size of the computational arena was chosen to contain all movements (3.34 *×* 1.67 *×* 0.4 cm in the forward, lateral and vertical directions respectively), with periodic boundary conditions imposed and the fish being initially positioned near the arena’s centre. The arena mesh was comprised of four adaptive layers, where the innermost layer had cell dimensions 32.7 *×* 32.7 *×* 16.34 *µ*m in the *x × y × z* directions. Fluid viscosity was set prior to each simulation at 0.00835 Pa.s for water and 0.0273 Pa.s for viscous simulations. Fluid velocity was initialized to zero.

### 3D model

To generate the 3D models, several fish at each age were fixed for 3 hours in 4% paraformaldehyde (PFA) in Phosphate buffered saline (PBS), washed in PBS with 0.1% Tween^®^ 20 (PBST), placed in a photo-bleach solution exposed to UV light four times (for 5 total minutes), then re-washed in PBST. 1:50 Phalloidin-Alexa 488 was then added to the PBST for 5 minutes. Fish were then washed for another 5 minutes in PBST, embedded in 1.2% agarose, and imaged on a Zeiss 710 confocal microscope using 488 laser at 256 X 256 using 10X objective, and a z-stack of 6*µ*m with approximately 50 levels.

Scans were filtered by a fixed pixel intensity that varied between fish, and the best scan from each age group was selected. The exterior layers were extracted, mirrored, and the two halves stitched together to create complete fish. Due to differences in embeddings between fish, manual adjustments were made prior to the mirroring and stitching. These included manually pushing caudal tail vertices in the x,y, and z directions. Manual adjustments were also made after the mirroring and stitching, due to differences in embedding, microscope penetration distances, and pixel intensities between fish scans. These involved cropping over-exposed eye regions, enlarging under-exposed nose regions, and matching top-down tail outlines with top-down reference pictures of larval zebrafish. Outlier vertices were removed via airbrushing.

To prepare the models for the CFD method, the models’ midlines were segmented into 125 evenly-spaced elliptical rings. Each elliptical ring was then uniformly filled by points. To reduce simulation time the 3D models were grid-desampled using a mesh distance of 30*µ*m. The models chosen had n=16,180, 16,491, and 18,496 vertices for 5, 9, and 14dpf respectively. Models were then scaled to match the respective lengths of each individual fish prior to simulation.

### CFD method

The CFD method was adapted from [54]. Briefly, for each time-step, we mapped the relative motion from experimentally-tracked swim bouts onto the 3D model and estimated point-wise velocities. To map relative motion, the relative x-y displacements of each midline point were directly mapped to all vertices in the corresponding elliptical slice of the 3D model. Relative z-displacements were set as 0 for all vertices. To estimate point-wise velocity, we computed the change in relative x-y displacements for the previous and future time-steps. Velocity at t=0 was set to 0. Point-wise velocities were then mapped to the respective elliptical slices. To prevent unrealistic deformations in the 3D model during turning movements, elliptical slices were rotated along the z-axis. The rotation magnitude corresponded to the change in local curvature between time-steps for the relevant midline point. To determine turning angle, two points on the 3D models were tracked during the simulation, one near the eye-midpoint and another near the swim bladder. CFD movies were created using VisIT [87].

Code for running the CFD simulations (once IBAMR has been locally installed) is provided at https://github.com/GoodhillLab/ZFLOW, along with an example of its application to a swim bout.

### CFD validation

To validate the CFD simulations, we first compared the mean hypotenuse velocity between each experimental swim bout (ground truth) and its simulated swim bout (IBAMR’s prediction), where Hypotenuse 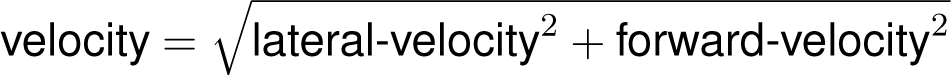. Velocities were computed using the eye-midpoint of the 2D tracking and the 3D models. We then computed the relative error (RE) as experimental hypotenuse velocity minus simulated hypotenuse velocity, divided by experimental hypotenuse velocity.

Several criteria were used to identify and exclude failed or inaccurate simulations. First, failed simulations were identified by either comparing the number of time-steps between a simulated bout and its original experimental recording; failed simulations were defined as having more than 10 time-step differences, or by producing NaNs in the output files. Failed simulations almost always occurred due to critical Courant–Friedrichs–Lewy (CFL) violations. Second, computationally-unstable simulations with non-critical CFL violations sometimes triggered adaptive time-steps within the CFD method that caused inaccurate velocity estimates. To identify these simulations, we compared the total time between experiment and simulation for each swim bout and removed simulations if the difference was larger than 5 microseconds. To ensure only accurate simulations were analysed, we removed simulations with RE*≥*0.5.

### CFD Energetics

The total energy cost (ergs) for each swim bout was calculated as *E_tot_* = ^J^ *|P_x_|* + ^J^ *|P_y_|*, where *P_x_* and *P_y_* are the power traces in the x and y-directions. Power in the z-direction was ignored because it had negligible size for the majority of simulations.

To estimate fish mass, 30 larvae were picked from each age group and euthanized using MSD22. The larvae were then placed in a weighing boat, and the excess water was carefully removed from the surface of the larvae. The total mass of the 30 larvae were measured using a fine scale balance. This procedure was repeated four times for 5 and 9 dpf fish and twice for 14 dpf fish.

### Measures of hunting efficiency

To estimate hunting efficiency, Capture Cost was defined as the total energy spent during capture events, divided by the number of captured *Paramecia*. Abort Cost was defined as the total energy spent during abort/miss events, divided by the total number of abort events. Event Cost was defined by the total energy spent during hunting, divided by the number of hunting events. Total Cost was defined as the total energy spent during all hunting events, divided by the total number of captured *Paramecia*.

### Kinematic measurements

For each swim bout, tail-beat amplitude of a bout was defined as the maximum tail-tip distance from the midline normalized by fish length. The midline was defined as the average y-position of fish midline at time=0. Tail-beat frequency was defined by the peak tail-tip frequency through time from a Fast Fourier Transform. For both measurements, to better account for turning motions, we removed net fish rotation and translation prior to measurement. Tail curvature was defined by computing angles between consecutive midline points and taking the mean for each time-step. Swim speed was defined by the absolute eye-midpoint displacement divided by bout duration.

Total kinetic energy was calculated by integrating the 0.5*mv*^2^ curve for each swim bout, where *m* is the mass of the fish and *v* is the hypotenuse velocity of the eye midpoint.

## Acknowlegements

The authors acknowledge the Research Infrastructure Services (RIS) group at Washington University in St. Louis for providing computational resources and services needed to generate the research results delivered within this paper. URL: https://ris.wustl.edu/

---

**Supplementary Figure 1:**
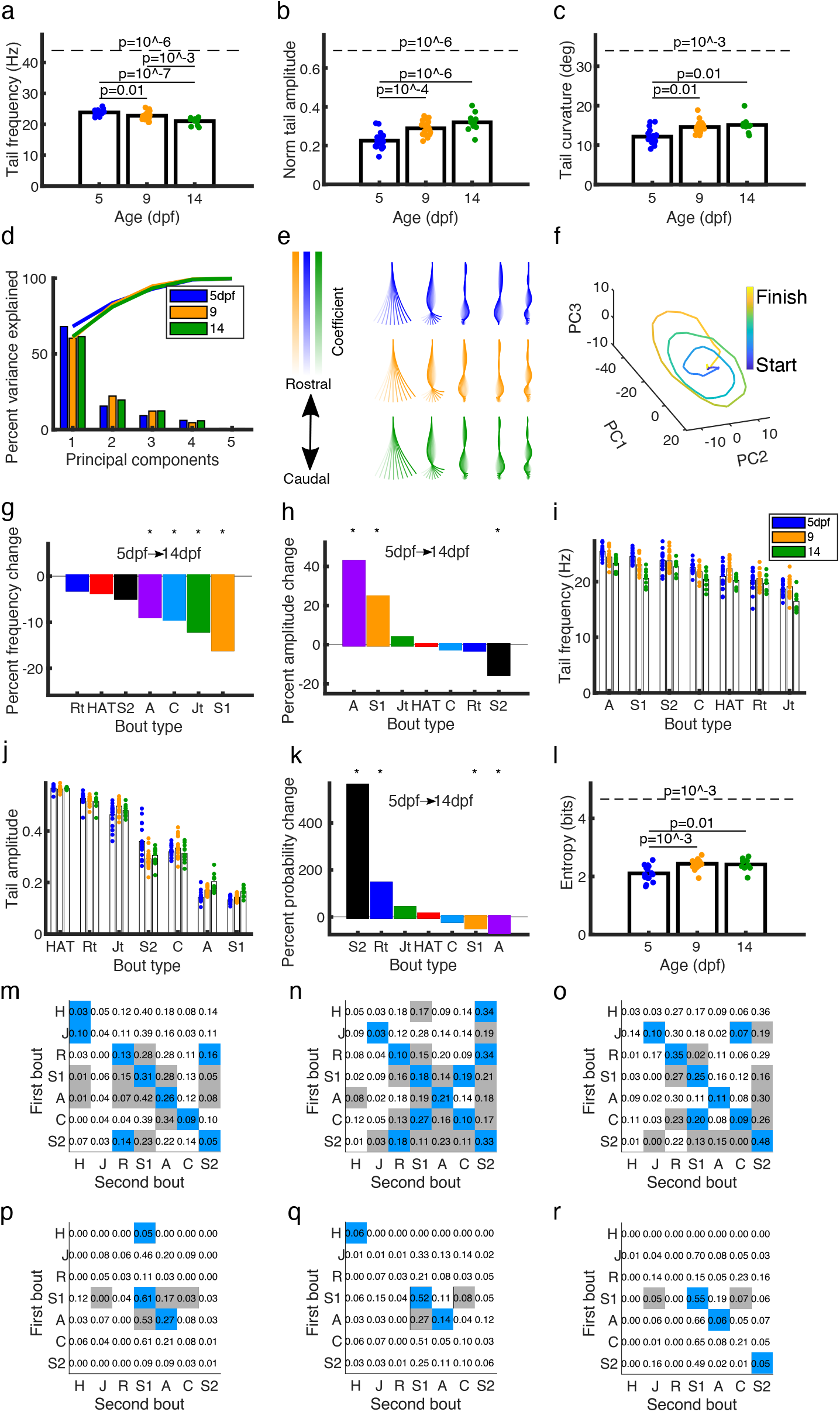
Changes in movement measures over development. **a-c** Tail-beat frequency decreased while tail-beat amplitude and curvature increased over development. Dashed p-value line indicates ANOVA, solid p-value lines indicate post-hoc pairwise comparisons. **d** There was little difference in variance explained by the leading PCs when calculated for each age separately. **e** There were no qualitative differences in the shape and ordering of eigenfish when calculated for each age separately. **f** An example PC trajectory of a 9 dpf Slow 1 bout. **g-h** Slow 1 and Approach bouts had changes in both tail-beat frequency and amplitude over development. Asterisks denote significant pairwise difference in means for a given bout type between 5 and 14 dpf. **i-j** Differences in frequency and amplitude between bout types were generally larger than changes within bout types over development. Each dot is the average from one fish. **k** There were significant changes in bout-type probabilities for Slow 2s, Routine turns, Slow 1s, and Approach bouts between 5 and 14 dpf. **l** There was an increase in entropy over development for the bout-type probability distributions. **m-r** Bout-type transition probabilities during exploring (**m-o**) and hunting (**p-r**) were relatively stable over development (5, 9, 14 dpf respectively). Blue, grey, and white squares represent transitions that occur more than expected, less than expected, and at chance level respectively (see Methods).

**Supplementary Figure 2:**
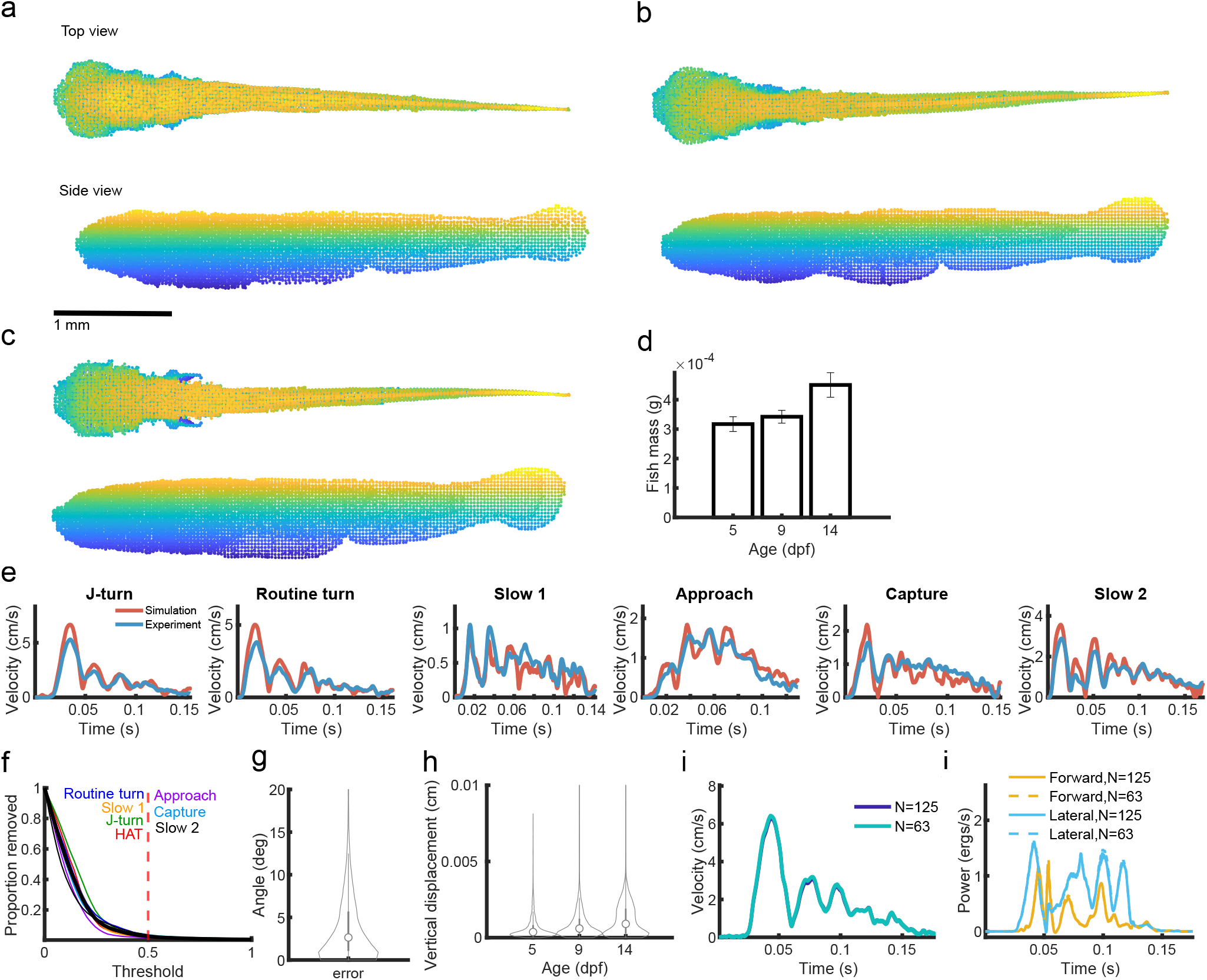
CFD simulation preprocessing and validation. **a-c** 3D models used for CFD simulations for 5, 9, and 14 dpf respectively (color represents height). **d** Fish mass increased over development. **e** There was a good match between simulated and experimental eye-midpoint velocities for all bouts types. The bouts shown here have REs reflecting the median for their respective bout type. **f** The proportion of bouts removed as a function of RE threshold. Colored lines are the proportion for each bout type, dashed red line is the chosen accuracy threshold of 0.5. **g** The distribution of simulation errors for angle. **h** Vertical displacements were very small. **i-j** Model predictions were insensitive to halving the number of tracking points.

**Supplementary Figure 3:**
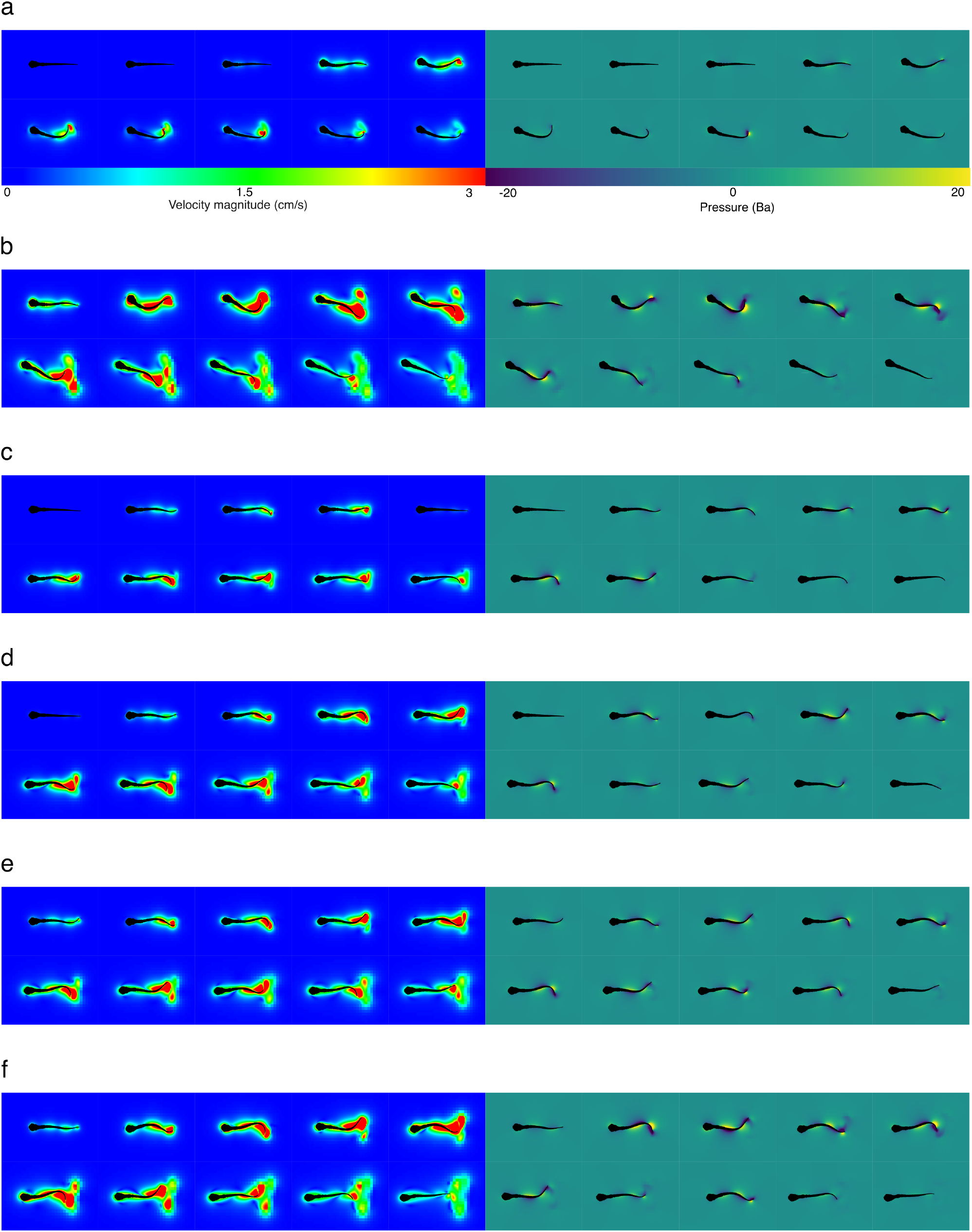
Visualization of example bout types. Example 2-dimensional snapshots of CFD simulations for each bout type, with *≈*10 ms spacing between frames. Left: The absolute velocity magnitude field. Right: The pressure field. **a** J-turn. **b** Routine turn. **c** Slow 1. **d** Slow 2. **e** Approach. **f** Capture Strike.

**Supplementary Figure 4:**
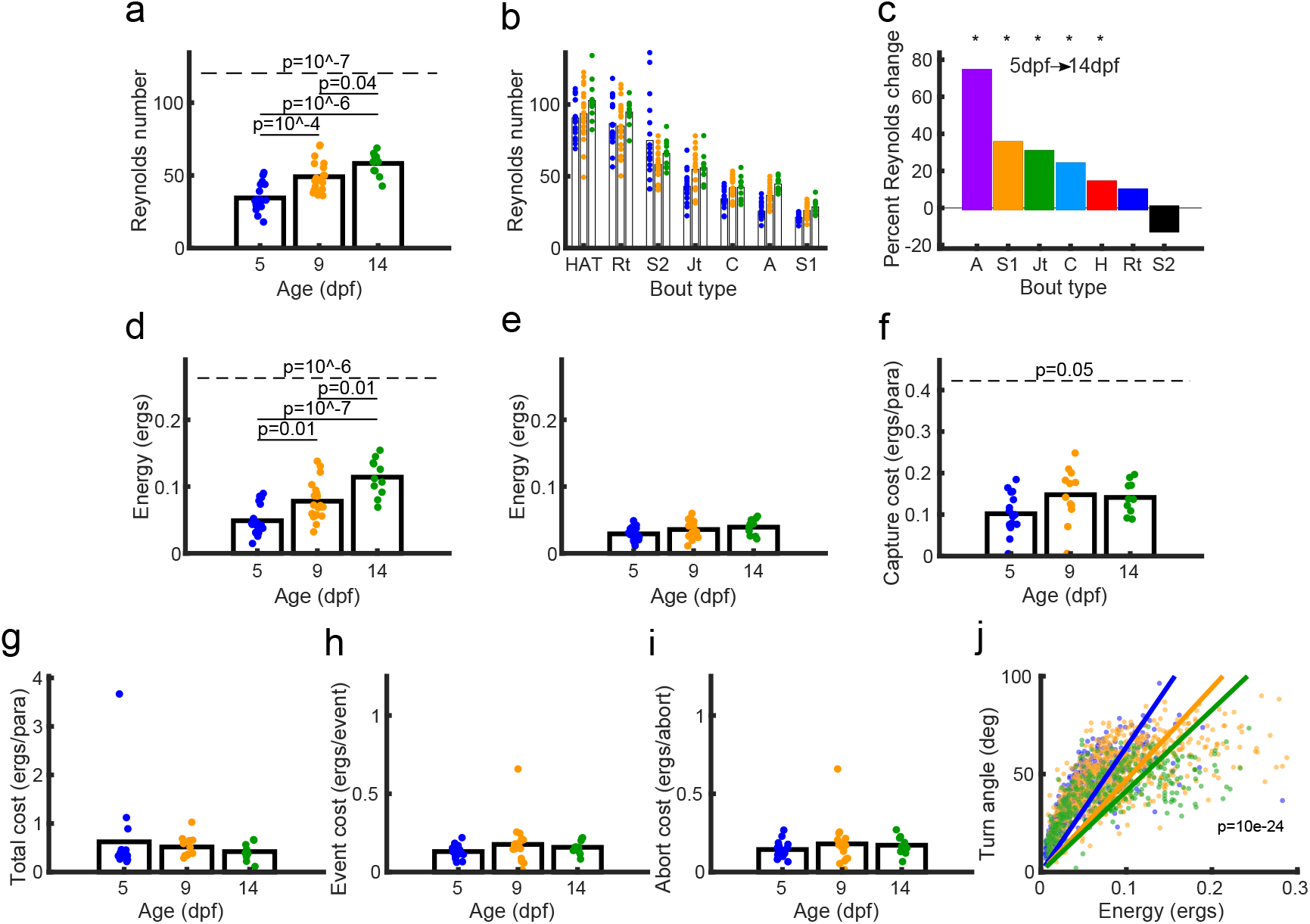
Energetics measures for larval zebrafish behavior. **a** Reynolds number increased over development. **b-c** This increase was dependent on bout type. Asterisks in c denote significant differences between 5 and 14 dpf. **d-e** There was a developmental increase in energy cost for exploring (d) but not for hunting (e). **f** There was an increase in Capture Cost over development. **g-i** There was a no change in Total Cost, Event Cost or Abort Cost over development. **j** For J-turns, the energy required to turn any specific angle increased with age. Dots represent individual bouts, colored lines represent fitted linear regressions *y* = *β*_1_*x* for each age group. Significance value derived from linear regression test for different slopes.

**Supplementary Figure 5:**
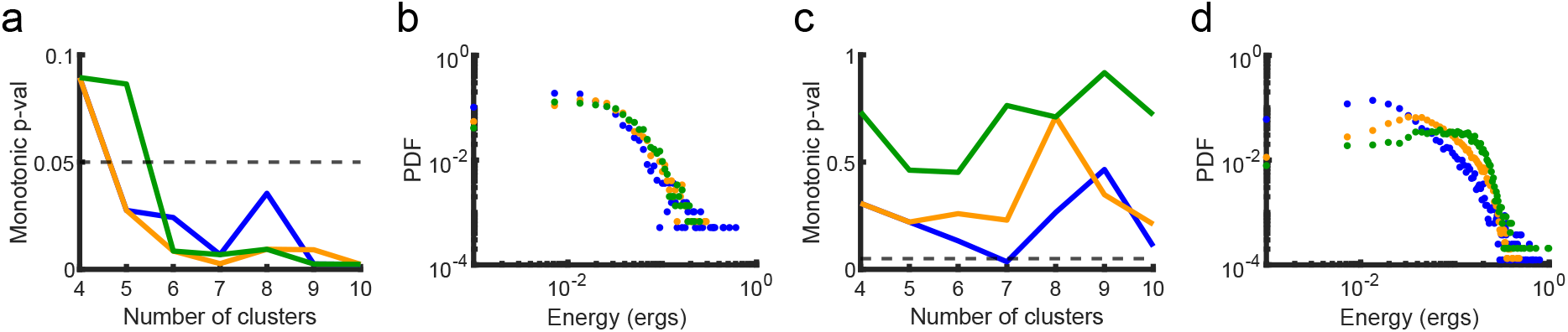
Monotonic relationship is robust against energy-binning methods. **a** The monotonic relationships between bout-type energy cost and probability for hunting were fairly stable across a clustering range of n=4 to n=10. **b** The monotonic relationships between bout-type energy cost and probability for hunting were maintained when binning energy instead of using clusters. **c-d** During exploration monotonic relationships were not observed across different numbers of clusters and binning.

**Supplementary Figure 6:**
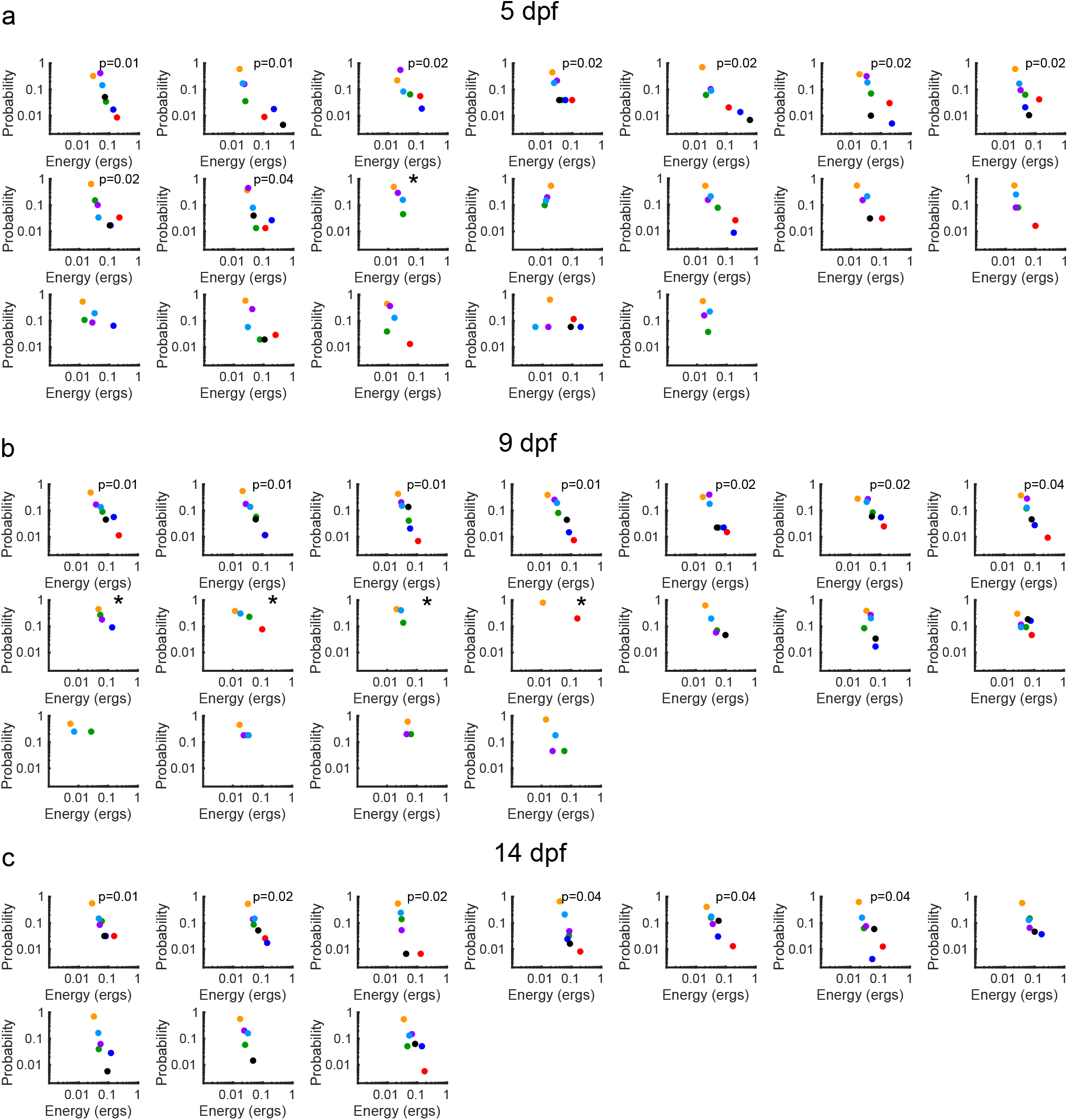
Monotonic relationship was present for the majority of individual fish. **a** 5dpf. **b** 9dpf. **c** 14dpf. Each plot is an individual fish, each dot is the bout-type probability and energy cost for that fish during hunting. Note that some fish use less than 7 bout types during hunting. *∗* is used to indicate a monotonic relationship in fish which used less than 5 bout types, since the Mann-Kendall monotonicity test is not reliable in this case.

**Supplementary Figure 7:**
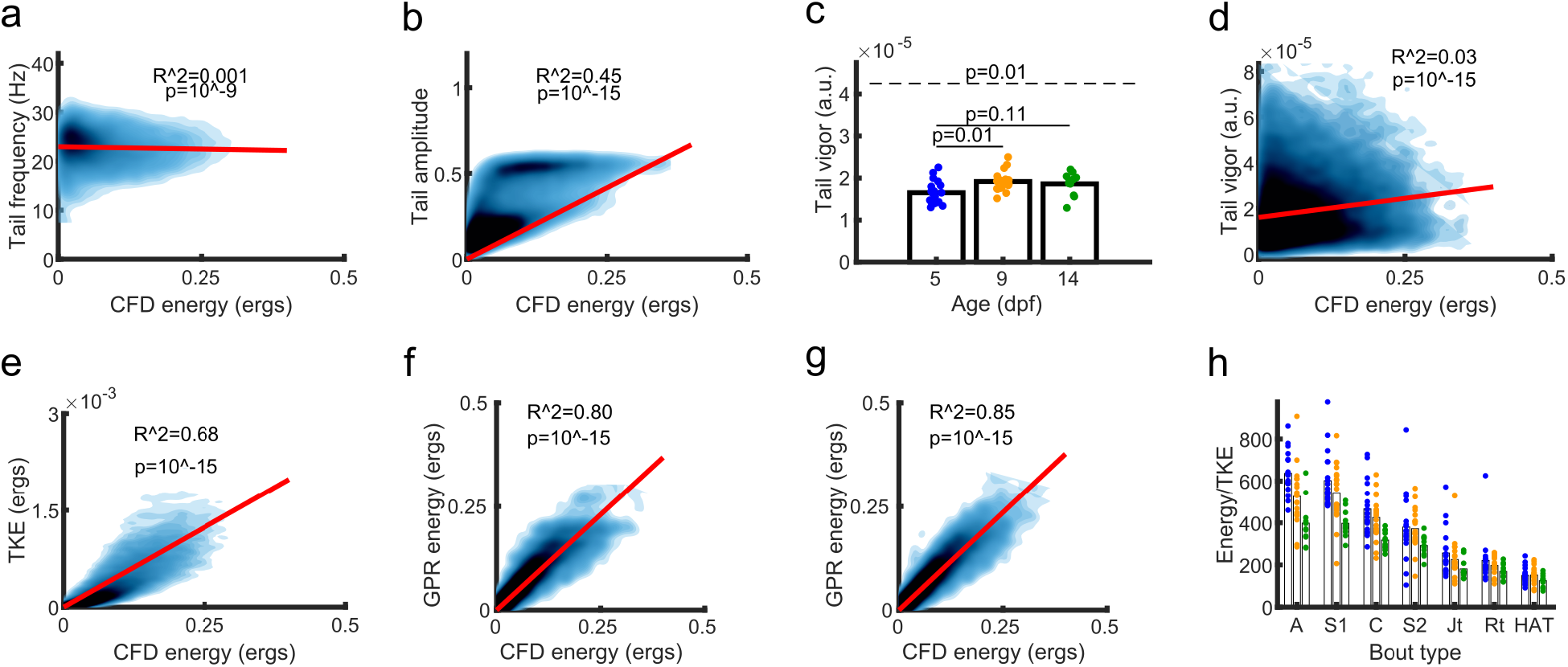
Approximations to CFD energy. **a** Tail-tip frequency has a very low correlation with energy usage. Red line represents a fitted linear regression *y* = *β*_0_ + *β*_1_*x*. **b** Tail-tip amplitude has a low correlation with energy usage. Red line represents regression *y* = *β*_0_ + *β*_1_*x*. **c** Tail vigor increases over development. **d** Tail vigor has a medium correlation with energy usage. Red line represents a linear regression line *y* = *β*_0_ + *β*_1_*x*. **e** TKE has a medium correlation with energy usage over development. Red line represents a fitted linear regression *y* = *β*_1_*x*, where *β*_1_ = 1/202. **f** A SVM trained on absolute displacement (*displacement_total_*) for swim bouts (see Methods) has a high correlation with CFD energy usage. Red line represents a linear regression line *y* = *β*_1_*x*. **g** A SVM trained on the final displacements (*displacement_x_, displacement_y_*) (see Methods) has a high correlation with CFD energy usage. Red line represents a linear regression line *y* = *β*_1_*x*. **h** The ratio of CFD energy to TKE differed with age and with bout type.

**Supplementary Figure 8:**
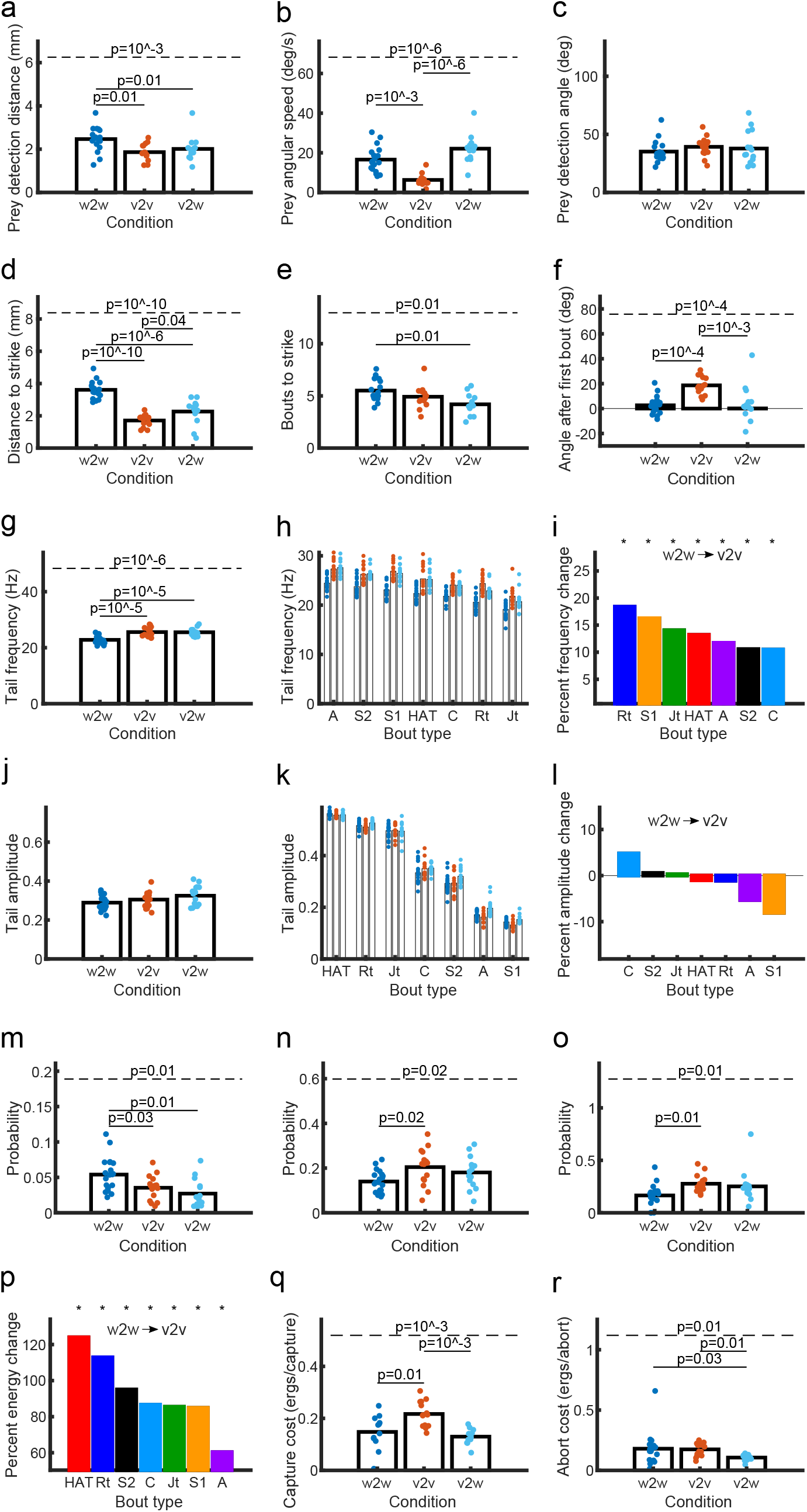
Viscous rearing affects hunting metrics but not movement mechanics. **a-b** The v2v fish hunted prey that were closer (a) and moving more slowly (b) than w2w fish. Dashed line has p-values from ANOVA, solid lines have p-values from pairwise comparisons, each dot is a fish average. **c** Prey detection angles were the same across conditions. **d** v2v fish travelled shorter distances during hunting events. **e** The number of bouts to strike was different between conditions. **f** v2v fish had larger undershoots during the initial hunting turn than w2w fish. **g** There was no difference in tail-beat frequency between v2w and w2w fish. **h** There were differences in tail-beat frequency between bout types. **i** There were differences in tail-beat frequency between v2v and w2w fish. **j** There was no difference in tail-beat amplitude between v2v and w2w fish. **k** There were differences in tail-beat amplitude between bout types. **l** There were no differences in tail-beat amplitude between v2v and w2w for all bout types. **m-n** There were differences in probability of J-turns (m) and Routine turns (n) during exploring between v2v and w2w fish. **o** There were differences in probability of Capture Strikes during hunting between v2v and w2w fish. **p** Increases in energy in the v2v compared to w2w case were dependent on bout type. **q** There was an increase in capture costs in v2v fish. **r** v2v fish use the same energy during abort as w2w fish.

**Supplementary Figure 9:**
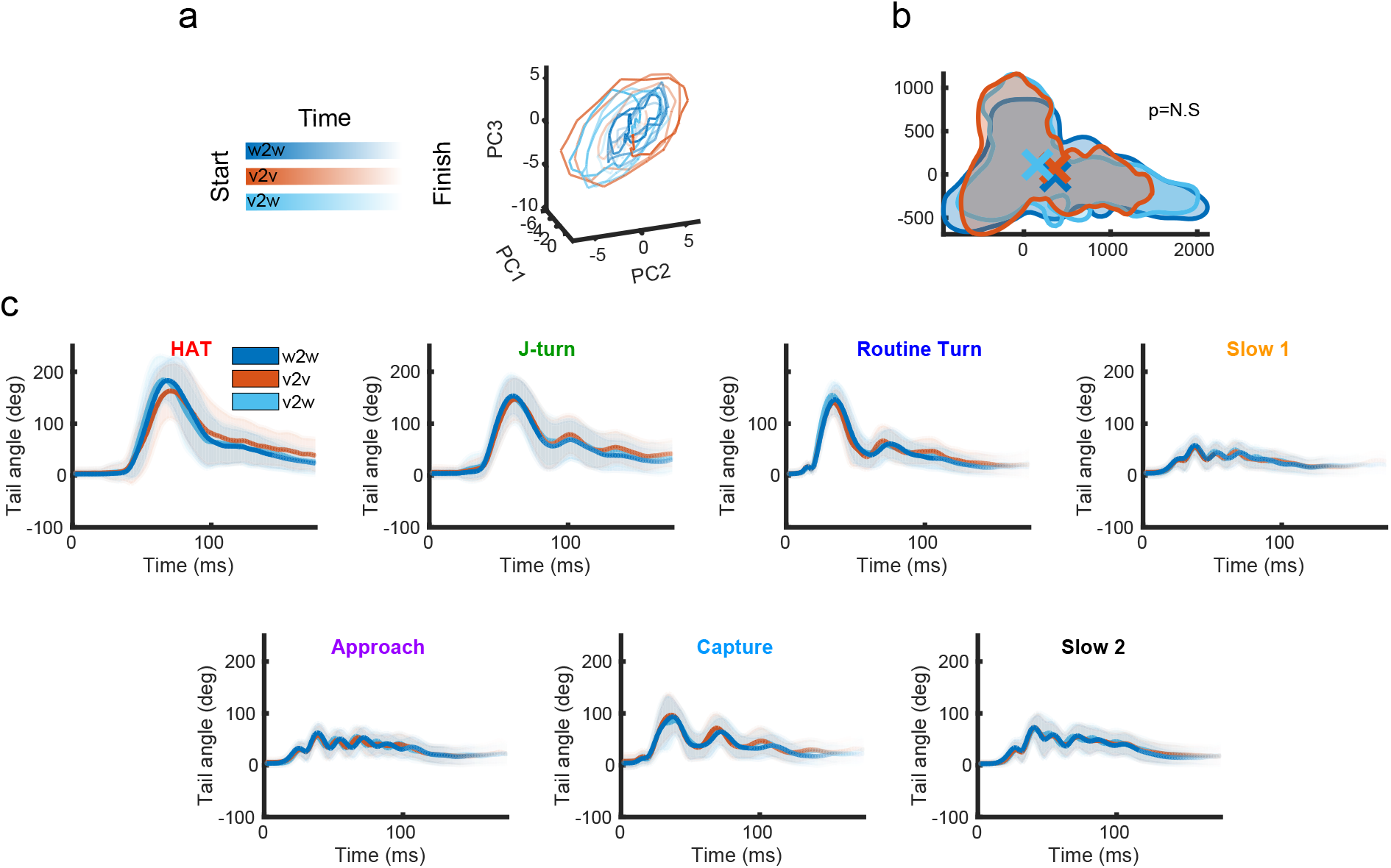
Viscous rearing does not affect low-dimensional structure of movement. **a** PCA trajectories of example Slow 1 swim bouts. **b** There was no difference in the areas covered by conditions within the combined Isomap embedding space. Shaded regions are outer layers of kernel density estimates for each condition, colored crosses are the center of mass for each condition in the embedding space. **c** For each bout type, there were no qualitative differences in the mean or variance of the tail-tip traces between the conditions. Thick lines indicate the average, shaded regions indicate 1 std.

**Supplementary Figure 10:**
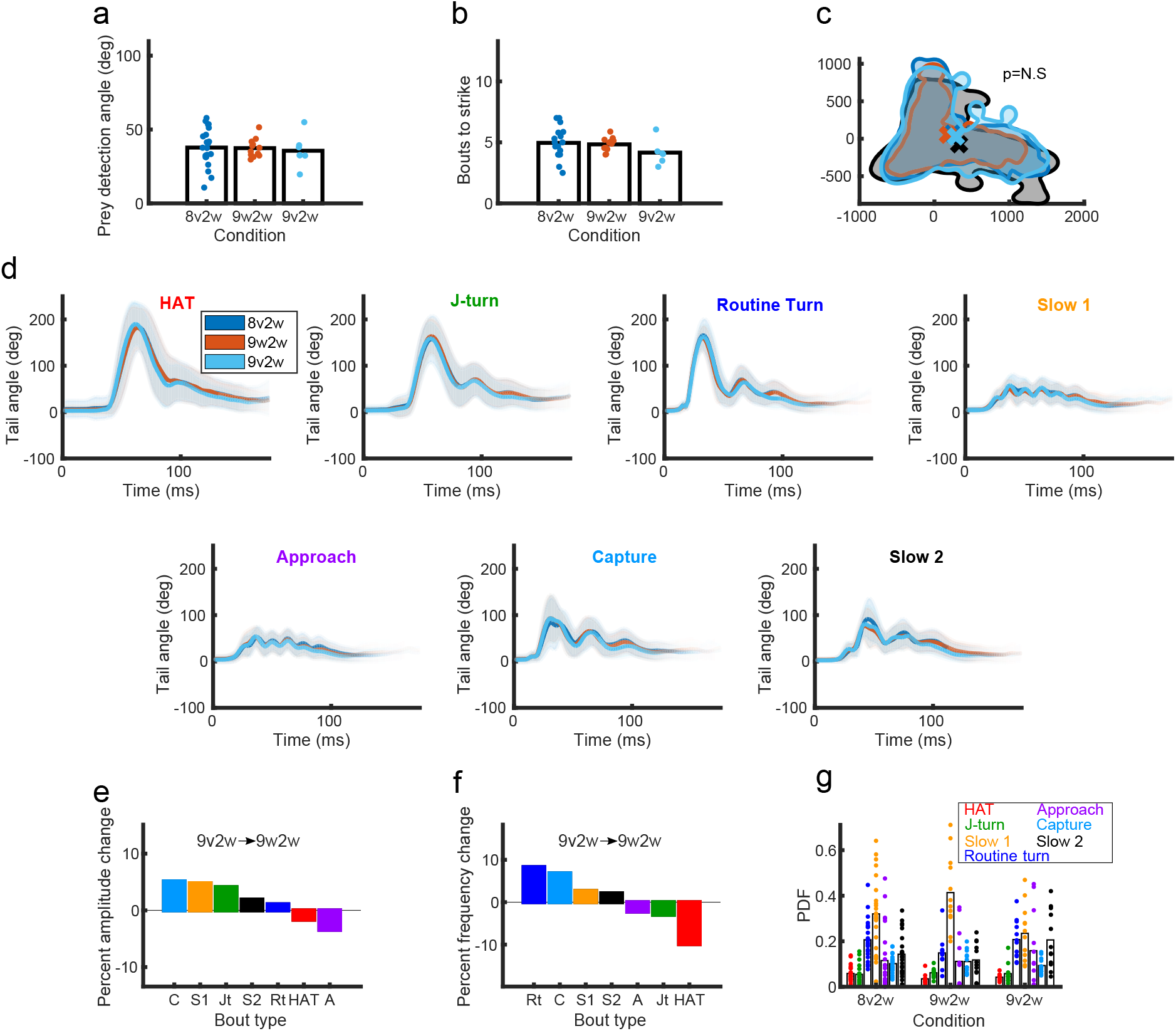
24 h adaptation to a novel fluid environment occurs primarily through bout-type probabilities. **a-b** Prey detection angle and bouts to strike and were similar between conditions. **c** There was no difference in the areas covered within a combined isomap embedding space. The shaded black region indicates 9dpf w2w exemplars from Fig. 1. **d** For each bout type the tail-tip traces were similar between conditions. Thick lines indicate average across all fish, shaded regions represent 1 std. **e** There were no significant differences in tail amplitude between 9v2w and 9w2w fish. **f** There were no significant differences in tail frequency between 9v2w and 9w2w fish. **g** There were differences in bout-type probabilities between conditions across all behaviors. Each dot represents a fish. For all possible comparisons between conditions, the p-values *<* 0.0001 (discrete KS-test).

